# Large-scale identification of phospho-modulated motif-based protein-protein interactions

**DOI:** 10.1101/2022.06.08.495335

**Authors:** Johanna Kliche, Dimitriya Hristoforova Garvanska, Leandro Simonetti, Dilip Badgujar, Doreen Dobritzsch, Jakob Nilsson, Norman Davey, Ylva Ivarsson

## Abstract

Phosphorylation is an extensively studied post-translation modification that regulates protein function by promoting, inhibiting or modulating protein-protein interactions. Deciphering which of the hundreds of thousands of phosphosites in the proteome that regulate interactions remains challenging. We generated a proteomic peptide-phage display (ProP-PD) library to screen for phosphosites that regulate short linear motif-based interactions. The phage peptidome covers 13,500 phospho-serine/threonine sites found in the intrinsically disordered regions of the human proteome, each phosphosite being represented as a wildtype and a phosphomimetic variant. We screened 73 modular domains and identified 252 putative phospho-modulated interactions. Affinity measurements confirmed the phosphomodulation of 16 out of 21 tested interactions. We discovered a novel phospho-dependent interaction between clathrin and the mitotic spindle protein hepatoma-upregulated protein (HURP). We validated the phospho-dependent clathrin interaction in a cellular context and found it to be essential for the mitotic function of HURP. Structural characterisation elucidated the molecular basis for the phospho-dependency of the clathrin-HURP interaction. Collectively, our work showcases the power of phosphomimetic ProP-PD to discover novel phospho-modulated SLiM-based interactions required for cellular function.

## 1. Introduction

Protein phosphorylation is an ubiquitous post-translational modification, which provides the cell with an effective mechanism to transmit information on the cell state encoded in kinase activity to the rest of the proteome (1,2). Phosphorylation occurs most frequently on serine, threonine and tyrosine residues and the addition of an acidic phosphate moiety can have significant impact on protein-protein interaction (PPI) networks (3). Phospho-modulated interactions play a critical role in various cellular processes, orchestrating metabolic and cellular fates, mediating signalling and transport, as well as participating in structural organisation (1,4). Large-scale mass spectrometry-based proteomic analysis has provided evidence for nearly 300,000 phosphosites (PhosphoSitePlus, Dec 2021) (5,6). There is thus compelling evidence for the abundance of phospho-modulated cellular functions. However, the functional relevance of the majority of identified phosphosites has been questioned (7) due to the off-target activity of kinases which may result in promiscuous phosphorylation (8). Identifying biologically relevant phosphosites and establishing their function is a major challenge. Several experimental and bioinformatical approaches have been developed to pinpoint key functional phosphosites, including time correlation between kinase activation and the event of phosphorylation (9), evolutionary conservation (10), and a combination of intrinsic features of phosphosites, such as disorder score and associated kinases (11). However, to functionally assess the molecular and cellular function of individual phosphosites requires time-consuming biophysical experiments, as well as their validation in cell biology.

Phosphorylation frequently regulates interactions mediated by short linear motifs (SLiMs), which are compact binding interfaces (typically 3-10 amino acid long stretches) found in the intrinsically disordered regions of the proteome (12,13). The effects of phosphorylation on binding can range broadly. Phosphosites located within, or in close proximity of, binding motifs can act as binary on/off switches or tune the affinity of the SLiM-based interactions (1). Numerous experimental approaches have been used to explore phospho-dependent interactions. For example, Tinti *et al*. applied pTyr-peptide chips in order to probe for the binding preference of SH2 domains (14). Following alternative two-hybrid approaches, Grossman *et al*., performed yeast-two-hybrid experiments with a co-expressed kinase to capture pTyr-dependent interactions (15) and Aboualizadeh *et al*., used a mammalian membrane two-hybrid system to explore the phospho-dependent interactome of the anaplastic lymphoma kinase (16). Moreover, hold-up assays have been used to screen for phospho-regulated interactions across the PDZ domain family (17). Nonetheless, there is still a paucity of large-scale methods available for sensitive and specific screening of phospho-regulated SLiM-based interactions.

Exploring phospho-modulated SLiM-based interactions poses a dual challenge, first identifying the SLiM-based interactions, and then subsequently, establishing the effect of phosphorylation on binding. To address the first challenge, we previously developed proteomic peptide-phage display (ProP-PD) (18–20). In ProP-PD, we computationally design a peptide library that tiles the disordered regions of a target proteome, and display them on the coat protein of the filamentous M13 phage. The phage peptidome is then used in selections against bait proteins, and the binding enriched phage pools are analysed by nextgeneration sequencing (NGS) to identify peptides specifically binding to the bait domain (21,22). The approach efficiently captures SLiM-based interactions on a proteome-wide scale (20).

To directly identify phospho-modulated SLiM-based interactions, we created a phosphomimetic ProP-PD library that tiles ~13,500 serine/threonine phosphosites found in the intrinsically disordered regions of the human proteome that were functionally prioritised based on proteomic, structural, evolutionary and regulatory evidence (23). The study builds on a previous small-scale pilot study on phosphomimetic ProP-PD where we used serine/threonine to glutamate mutations to mimic phospho-serine/threonine in 4,828 unique C-terminal peptides (24). Here, we evaluated the general applicability of the approach by exploring the potential phospho-modulation of SLiM-based interactions of 73 domains from different classes. We identified 1,070 high-confidence phosphosite interactions for 52 domains, of which 24% were suggested to potentially be modulated by phosphorylation. We show the quality of the data by validating 16 of 21 (76%) of the discovered high-confidence phospho-modulated interactions. Among the novel phospho-modulated interactions, we highlight the interaction between the clathrin N-terminal domain (NTD) and the mitotic spindle protein Hepatoma-upregulated protein (HURP). Phosphomimetic ProP-PD thus allows the identification of phospho-modulated interactions with biological function.

## 2. Results

### 2.1 Design and quality of the internal phosphomimetic ProP-PD library

We designed a novel phosphomimetic human disorderome library (PM_HD2; **Table S1**) by combining the information on functionally prioritised phospho-serine/threonine-sites (23) with the design of our recently described second generation human disorderome (HD2) library (**Figure 1A, B**). The HD2 library tiles the disordered regions of the proteome using 16 amino acid peptides with 12 amino acids overlap between flanking peptides (20). The functional phosphosite score defined by Ochoa et al. (23) predicts functionally relevant phosphosites with a high probability to have an effect on phospho-modulated PPIs. The PM_HD2 design contains both the wild-type peptides with known serine/threonine phosphosites, and serine/threonine to glutamate (phosphomimetic) mutants thereof. This allows screening of the binding preferences of a given protein bait domain to either of the two peptide versions (wild-type or phosphomimetic) in a single experiment. A custom oligonucleotide library encoding the designed peptide sequences was obtained and was used to fuse the designed peptide sequences N-terminally to the major coat protein p8 of the M13 bacteriophage **(Figure 1A)**. NGS analysis of the constructed PM_HD2 library confirmed at least 91% of the designed sequences were present in the phage library, without any major sequence bias (**Figure 1C** and **Figure S1**).

**Figure 1.**
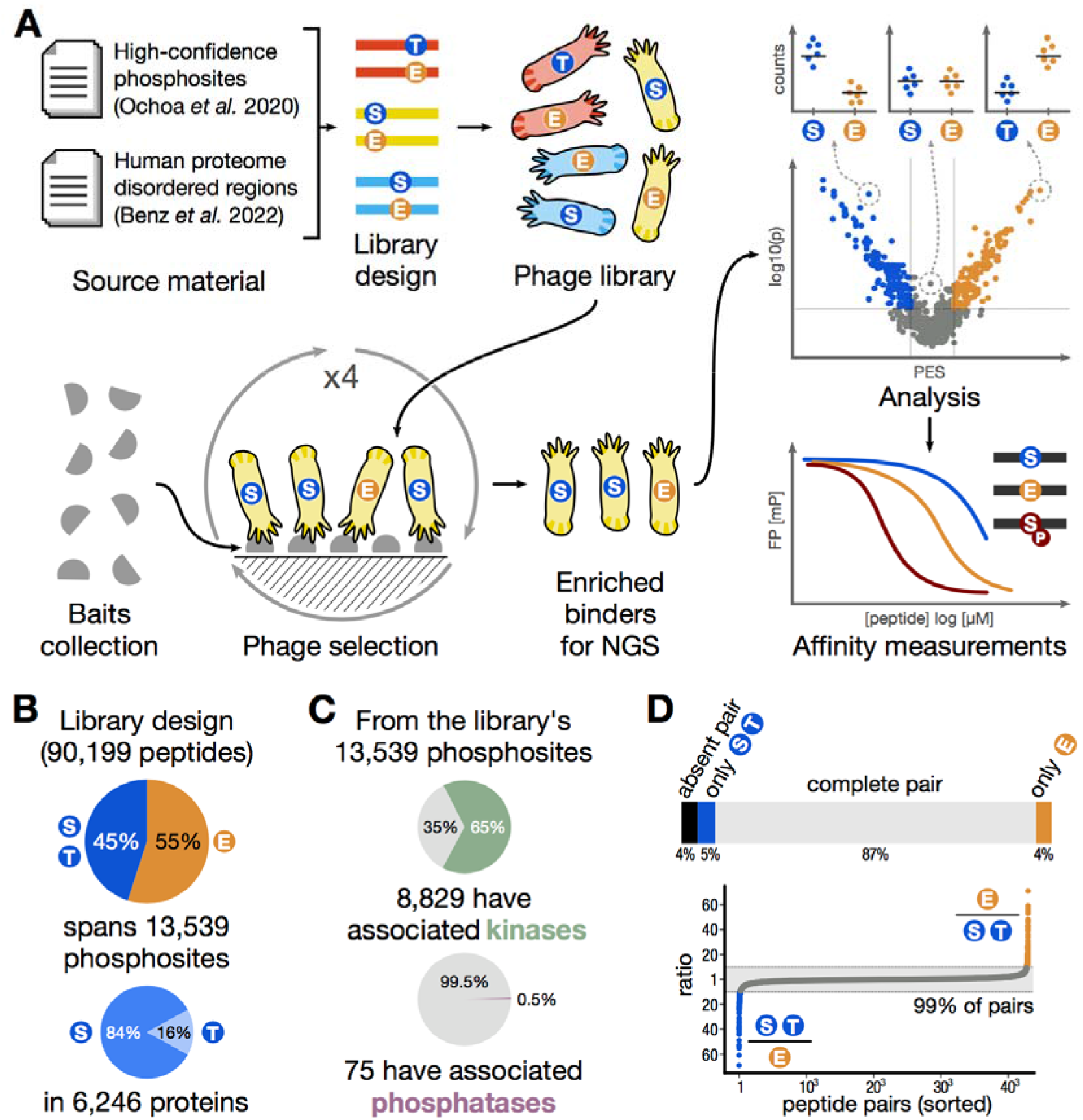
Overview of the workflow and details of the PM_HD2 library. **A:** Schematic overview of the workflow from library design and phage library construction, to data analysis and validations. **B:** Parameters of the PM_HD2 library design. **C:** Known associated kinases and phosphatases for the phosphosites included in the PM_HD2 library design. **D:** Completeness and proportions of the wild-type/phosphomimetic peptide pairs in the PM_HD2 library sequencing results.

### 2.2 The bait domain collection

To evaluate the ability of phosphomimetic ProP-PD to find phospho-modulated interactions, we designed a bait protein domain collection to cover well-known obligate phospho-peptide-binding domains (i.e. 14-3-3 proteins, PLK1 polo-box domain, IRF3 FHA domain and PIN1 WW domain), peptide-binding domains for which a phospho-modulation of binding was previously reported (26 domains), well-known peptide-binding domains used in our previous selections against the HD2 library (9 domains) (20) and additional peptide-binding domains to increase the diversity of interactions tested (**Table S2**). The bait proteins were glutathione transferase(GST)- or maltose binding protein (MBP)-tagged, and we therefore used GST and MBP as negative controls (**Figure 2A, B**). The domain collection encompassed 73 domains covering a variety of domain types (**Table S2**). The bait protein domains were used in four rounds of selections against the PM_HD2 library. Selections were successful for 52 out of 73 bait proteins tested based on phage pool ELISA analysis of the binding enriched phage pools. Notably, selections against obligate phospho-serine/threonine-binding domains largely failed (3 out of 10 successful), whereas the domains with previously reported effect of phosphorylation on binding (23 out of 26 successful) and the other peptide-binding domains (8 out of 9 successful) enriched for one or more medium/high confidence peptide during the selections (**Figure 1C**).

**Figure 2.**
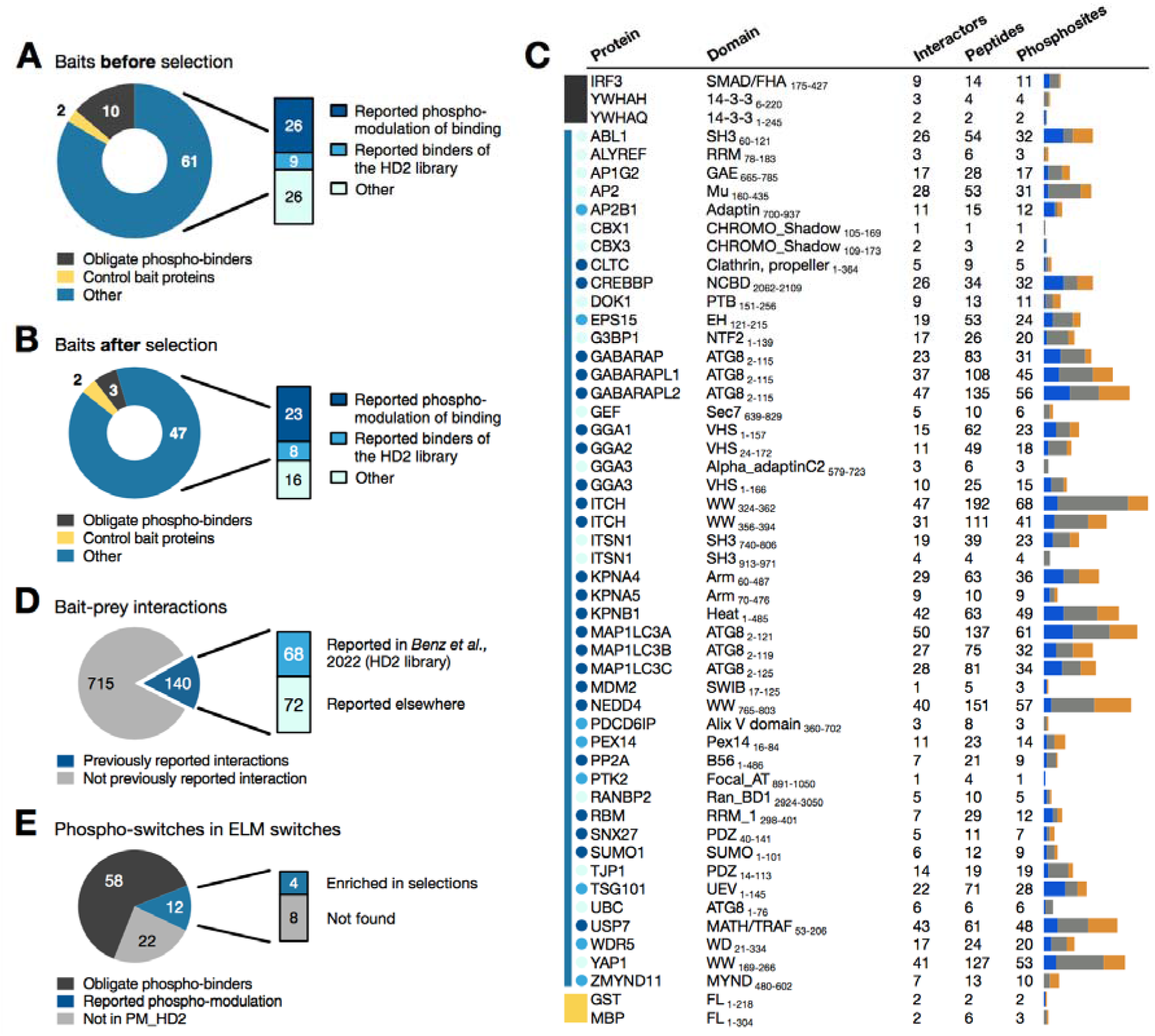
Baits collection and ProP-PD selection results. **A:** Overview of the bait protein collection screened against the PM_HD2 library. **B:** Bait domains which returned medium/high-confidence peptides in the screen against the PM_HD2 library. **C:** Selection results for each bait protein, including the total number of medium-/high-confidence peptides found and how many proteins and phosphosites do those peptides cover. The colours to the left of the gene names match those of panel A and B. The bars to the right show the number and distribution of phosphosites, for which we found evidence of phospho-modulation favouring the wild-type (blue), favouring the phosphomimetic (orange) or with no evidence of phospho-modulation found (grey). **D:** Percentage of interactions found that are supported by previous evidences by screening against the HD2 library or reported in HuRi and BioPlex databases (**Table S6B**)**. E:** Percentage of phospho-switches reported in switches.ELM (25) for the bait domain collection and the ones retrieved in selections.

### 2.3 Phage selection outcome and interaction analysis

The peptide-encoding regions of the enriched phage pools were barcoded and sequenced by NGS. The NGS results were analysed using a bioinformatic pipeline, where the sequences are translated to peptides, and then matched to the proteome using the PepTools peptide annotation tool (20). Peptides were assigned different confidence levels (confidence levels: 0 = no confidence, 1 = low, 2-3 = medium, and 4 = high confidence) based on an established set of criteria, including occurrence in replicate selections, identification of overlapping peptides and the presence of consensus motifs. For a stringent analysis, we here focus on peptide ligands with medium/high confidence (**Table S3**). In total, we found 855 PPIs involving 52 bait proteins and 512 unique putative phospho-proteins (preys), covering 681 unique phosphosites (for a total of 1,070 phospho-modulated interactions) (**Table S6**). Of the 855 interactions found, 140 (16%) were previously reported interactors, of which ca. 50% were found by screening with the HD2 library (20) and the remainder by other methods (**Figure 2D, Table S6B**).The switches.ELM resource (25) contains 92 previously reported phospho-switches for our bait domain collection of which 70 were part of the PM_HD2 design (**Figure 2E**, **Table S7**), however, most (58) relate to the obligate phospho-binders for which the selections returned no peptides or low confidence peptides. From the remaining 12 phospho-switches, 4 were found enriched in our selection experiments (**Figure 2E**). Among those is the binding of the MDM2 SWIB domain to p53, which is reduced by T18 phosphorylation (26).

The results for obligate phospho-binders contained no or few high confidence ligands, suggesting that the approach failed to identify ligands for these proteins. Nevertheless, we tested the binding of two of the obligate phospho-serine/threonine-binding domains (PIN1 WW domain and 14-3-3 theta) to putative binding peptides for which the display indicated a preference for the phosphomimetic peptide. We used wild-type, phosphomimetic and phosphorylated versions of the peptides in affinity determinations through fluorescence polarisation (FP) displacement experiment. The PIN1 WW domain bound none of the peptides tested (wild-type, phosphomimetic or phosphorylated PIAS2_493-507_). Similarly, 14-3-3 theta showed no binding for the wild-type and phosphomimetic peptide (TOM1_366-382_) and only weak affinity for the phosphorylated peptide tested (**Figure S2**). In conclusion, we found that phosphomimetic ProP-PD does not capture ligands of the tested obligate phosphobinding domains, possibly due to a strict requirement of a phosphate moiety rather than the substituted glutamate for binding to the obligate phosphor-binders in the domain collection. The remainder of our study focussed on cases in which phosphorylation abolished or modulated the affinity of interactions.

### 2.4 Identification of putative phospho-modulated SLiMs by screening with a phosphomimetic ProP-PD library

In order to deduce information on statistically significant mutation-modulated interactions from the phosphomimetic ProP-PD data, we analysed the normalised NGS sequencing counts to calculate a phosphomimetic enrichment score (PES) for each wild-type and phosphomimetic pair (**Table S4**). This allowed us to score the observed effects of the phosphomimetic mutations in the ProP-PD data on the level of the different phosphosites by combining data from multiple peptides and replicates (**Figure 3A**). In order to attribute statistical power to the PES as our scoring metric, we calculate the p-value for each of the interactions based on the Mann-Whitney Test (**Figure 3A**). By plotting the p-values against the calculated PES, a scatter plot was obtained in which the left arm (blue) indicates interactions inhibited by the phosphomimetic mutation and the right arm (orange) indicates interactions enabled by the phosphomimetic mutation (**Figure 3A**). Using a p-value of 0.01 and an absolute PES >2 as cut-off, 252 out of 1,070 phosphosite-interactions (24%) interactions were suggested to be modulated by the phosphomimetic mutation, of which 50% were disabling and 50% were enabling.

**Figure 3.**
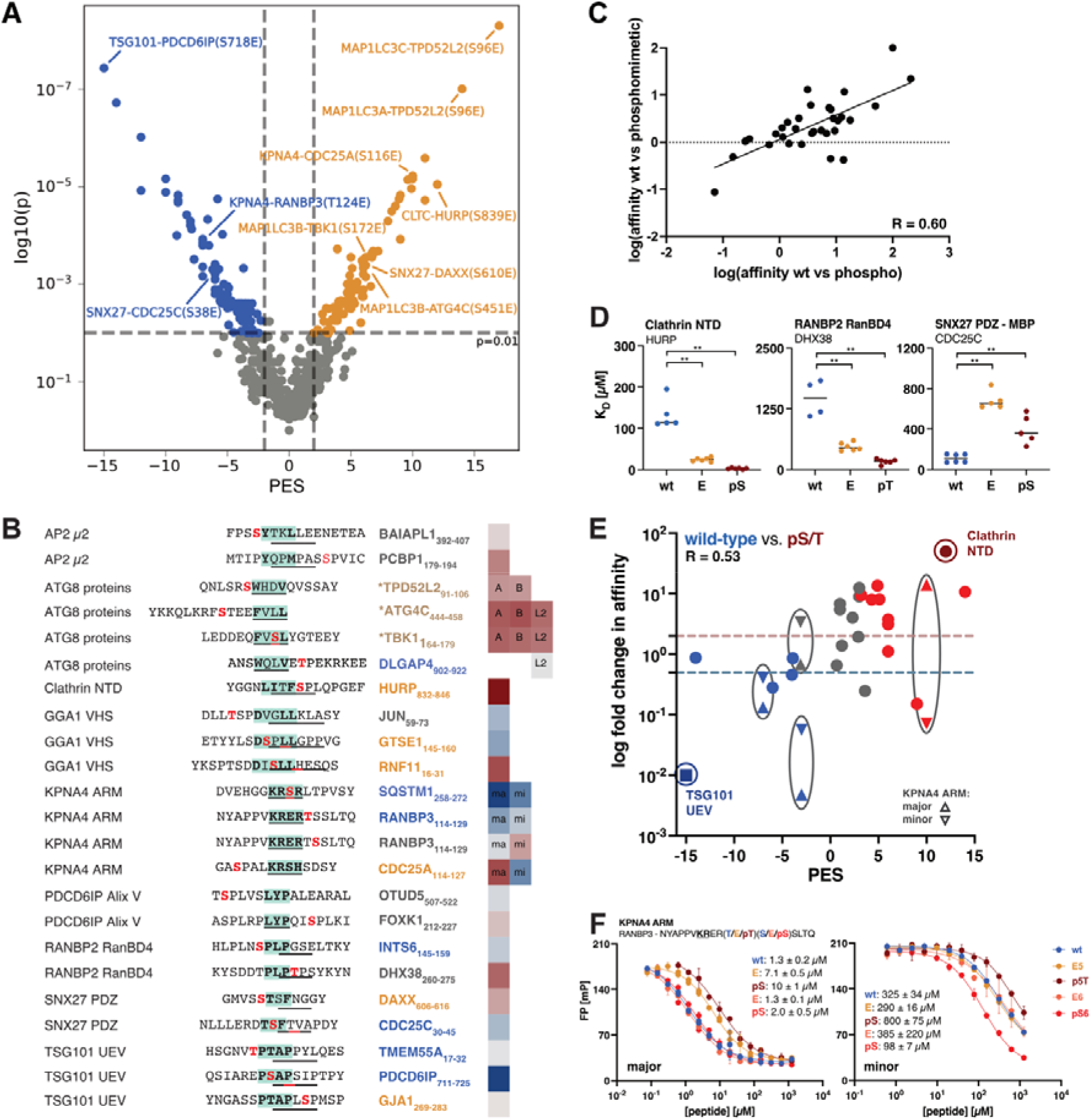
The phosphomimetic mutation in phage display serves as a good proxy for preferences for wild-type or phosphorylated peptides. **A:** Scatter plot of the PM_HD2 selection indicating pairs with a negative PES (left arm; preference for wild-type peptides) and positive PES (right arm; preference for the phosphomimetic peptides). Coloured peptides pairs represent cases in which the binding preference was statistically significant (Mann-Whitney-Test p ≤0.01 and an absolute PES value > 2) with blue pinpointing negative and orange pinpointing positive impact of the phosphomimetic mutation on the binding. **B:** Bait protein domains and peptides chosen for affinity measurements. In the peptides, binding motifs are indicated in green and underscored and the phosphosite in red. Affinity measurements were performed with the wild-type, phosphomimetic and phosphorylated version of the peptides. Coloured gene names of the prey proteins indicate phosphomodulated interactions of the display with p≤0.01 and absolute PES >2 (blue: disabling and red: enabling interactions). The measured fold change in affinity between the wild-type and phosphorylated peptide is indicated in blue (wild-type peptide with higher affinity) or red (phosphorylated peptide with higher affinity). A = MAP1LC3A; B = MAP1LC3B; L2 = GABARAPL2; ma = major, mi = minor. * = all ATG8 proteins have their individual PES value for those peptides. **C:** Correlation of the fold changes in affinity (K_D_ values) between wild-type/phosphomimetic and wild-type/phosphorylated peptide pairs (Spearman correlation R-value = 0.63). **D**: Examples of K_D_ values of wild-type, mutant and phosphorylated peptides binding to clathrin NTD (HURP_832-846_), RANBP2 RanBD4 (DHX38_260-275_) and SNX27 PDZ (CDC25C_30-45_). Significant differences indicated by the Mann-Whitney Test (**: p<0.01). **E:** Correlation between the fold change in affinity (wild-type vs. phospho-peptide) and the PES of the representative peptide pairs in the phage screen (Spearman R-value = 0.53). Red indicates positive and blue indicates negative phospho-modulation of the interaction. Grey colours indicate interactions outside the cut-off values p>0.01 and PES between −2 and 2. The dotted lines represent the log(2-fold change in affinity) cut-offs for enabling and disabling interactions respectively. **F**: Displacement curves of the KPNA4 ARM domain with the RanBP3_114-129_ peptides (wild-type, phosphomimetic and phosphorylated) sampling the effects of two distinct phosphosites (T124 and S125). IC50-values for the different peptides are indicated in μM.

We next assessed to what extent the phosphomimetic ProP-PD correctly captured phospho-modulated interactions. For this purpose, we addressed two core questions: (i) is the phosphomimetic glutamate mutation a valid proxy for phospho-serine/threonine, and (ii) does the PES from the phage display analysis capture the affinity differences between wildtype and phosphomimetic/phospho-peptides. We chose 32 putatively phospho-modulated interactions covering a window of the PES from −15 to 12 (**Figure 3A**) and determined affinities of the corresponding bait protein domains for wild-type, phosphorylated and phosphomimetic variants of peptides (in total 32 peptide triplets) through a FP displacement assay (**Figure S3**). The affinities were found to be in the low micromolar to millimolar range (**Table 1**).

**Table 1:**
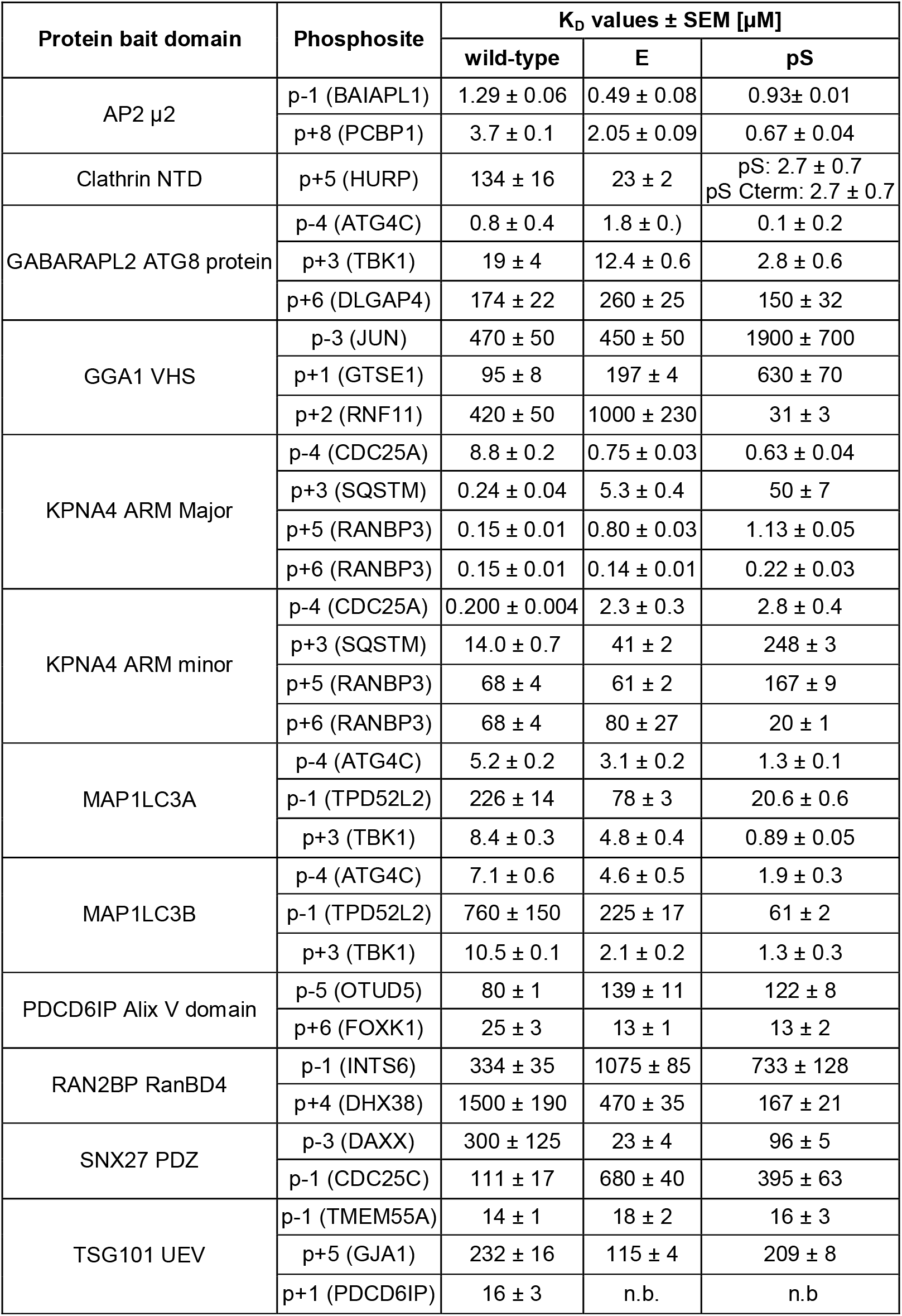
Protein bait domain, protein from which the FP peptides are derived, as well as the KD values of wild-type, phosphomimetic and phosphorylated peptides with SEM.

The comparison between the effect of phosphorylation and phosphomimetic mutation on binding (assessed by the fold change in affinity) revealed that there was a clear correlation, with phosphorylation conferring a more pronounced change in affinity (2-fold difference) than the phosphomimetic mutations (**Figure 3C**). This observation is likely explained by the net charge difference between a phospho-serine/threonine and glutamate. The interaction of SNX27 PDZ domain with ligands with an internal PDZ binding motif (xT/SxFx, with F = p0; found in DAXX6_06-616_ and CDC25C_30-45_) stood out as an exception to the general rule. The phosphomimetic mutations caused in both cases a higher fold-change in affinity in comparison to the phosphorylated peptides (p-3 phosphosite (DAXX6_06-616_): phospho/wild-type: 3-fold, phosphomimetic/wild-type; 13-fold; p-1 phosphosite (CDC25C_30-45_): wild-type/phospho: 3.5-fold; wild-type/phosphomimetic: 6-fold) (**Figure 3D, Table 1**). In order to strengthen our affinity data set, we repeated a selected set of FP measurements using independently prepared protein domains (**Figure 3D**) and complemented the affinity results by isothermal titration calorimetry (ITC) (**Figure S4, Table S7**), which gave further confidence in the affinity data. The FP repeat measurements confirmed significant differences in affinity for the wild-type peptides as compared to phosphomimetic and phosphorylated peptides for clathrin NTD for HURP peptides (enabling effect), RANBP2 RanBD4 for DHX38 peptides (enabling effect) and SNX27 PDZ for CDC25C peptides (disabling effect), consistent with the phosphomimetic ProP-PD results. In addition, affinities obtained by ITC measurements were in good agreement with the FP measurements (**Table 1, Table S7**, e.g. MAP1LC3A ATG8 protein: K_D_ = 8 μM (FP, TBK1 wild-type peptide) and K_D_ = 7 μM (ITC, TBK1 wild-type peptide).

We next evaluated, on the phosphosite level, if the calculated PES (−2 ≥ PES ≥ 2) and the associated p-value (p≤0.01) correlated with the fold change in affinity elicited by phosphorylation. For 16 of the 21 (77%) tested putatively phospho-regulated interactions there was a good agreement between the PES and the fold change in affinity (at least 2-fold) between wild-type/phosphorylated peptides in terms of effect on binding (**Figure 3E**). For 11 of the interactions tested the PES was between −2 and 2 and the associated p-value > 0.01, and for these cases phosphorylation had either no effect on binding or there having an effect with poor correlation with the PES. This argues in favour of the chosen cutoff values.

The PES performed similarly well in indicating disabling (7 out of 9 interactions) and enabling (9 out of 12 interactions) effects on binding. For 23% (5) of the cases, the PES failed to correctly predict the effect of phosphorylation on binding. For the binding of GGA1 VHS domain to the GTSE1 peptide, it can be argued that the affinities for all peptides was low, which renders the fitting of the data less accurate. Our validation set further included ligands for KPNA4, for which there is an added level of complexity in that the protein has two binding pockets (major and minor) that bind basic motifs of different classes (20). Using probe peptides for the distinct binding pockets we deciphered that the effects of the modifications were different depending on the binding pocket. Notably, we found that the PES reflected the change in affinity for the major pocket, suggesting that this was the main pocket probed during the phage selection. This becomes particularly apparent for the CDC25A_114-127_ peptides, for which the two binding pockets show opposite preferences for pS116 (**Table 1**). Overall, the analysis supports that the phosphomimetic mutation is a valid proxy for serine/threonine phosphorylation and tool for probing the effects of phosphomodulation on interactions.

### 2.5 Position-wise analysis of the effect of phosphomimetics and phosphorylation in relation to consensus motifs

Phosphomimetic ProP-PD offers the possibility to simultaneously identify SLiM-based interactions and to gain position-wise information on the effect of phosphosites in relation to the consensus motif of a given protein domain. A relevant question is whether it is possible to predict the effect of phosphorylation based on the results and motifs generated using a conventional ProP-PD library displaying only wild-type sequences or using a combinatorial peptide phage library. We hypothesise that the binding motifs of the individual domains have partial predictive value when it comes to position-wise phosphorylation with acidic residues serving as positive and serine/threonine residues as negative indicators and explored it using the MAP1LC3A ATG8 protein, as well as the TSG101 UEV and SNX27 PDZ domain. The ATG8 proteins are known to bind a LC3-interacting region (LIR) motif, which centres around an aromatic residue at p+1 position and a hydrophobic residue at p+4, interspaced by two variable amino acids ([F/Y/W]xx□]) (27,28) (**Figure 4A**). We tested the phospho-modulation of this interaction at position p-4, p-1 and p+3 for the MAP1LC3A ATG8 protein. The protein bound all tested phosphorylated peptides with higher affinity as compared to the wild-type peptides (**Figure 4B, C**, Table 1). A comparison to the residues favoured at these positions as determined by previous ProP-PD selections against the HD2 library ((20), **Figure 4A**), revealed a partial enrichment of acidic residues (aspartate/glutamate) at the p-4 and p-1 positions consistent with the observed effects of phosphorylation. In contrast, the positive effect observed for phosphorylation of the p+3 position was not revealed by the consensus motif, but by using the phosphomimetic ProP-PD approach.

**Figure 4.**
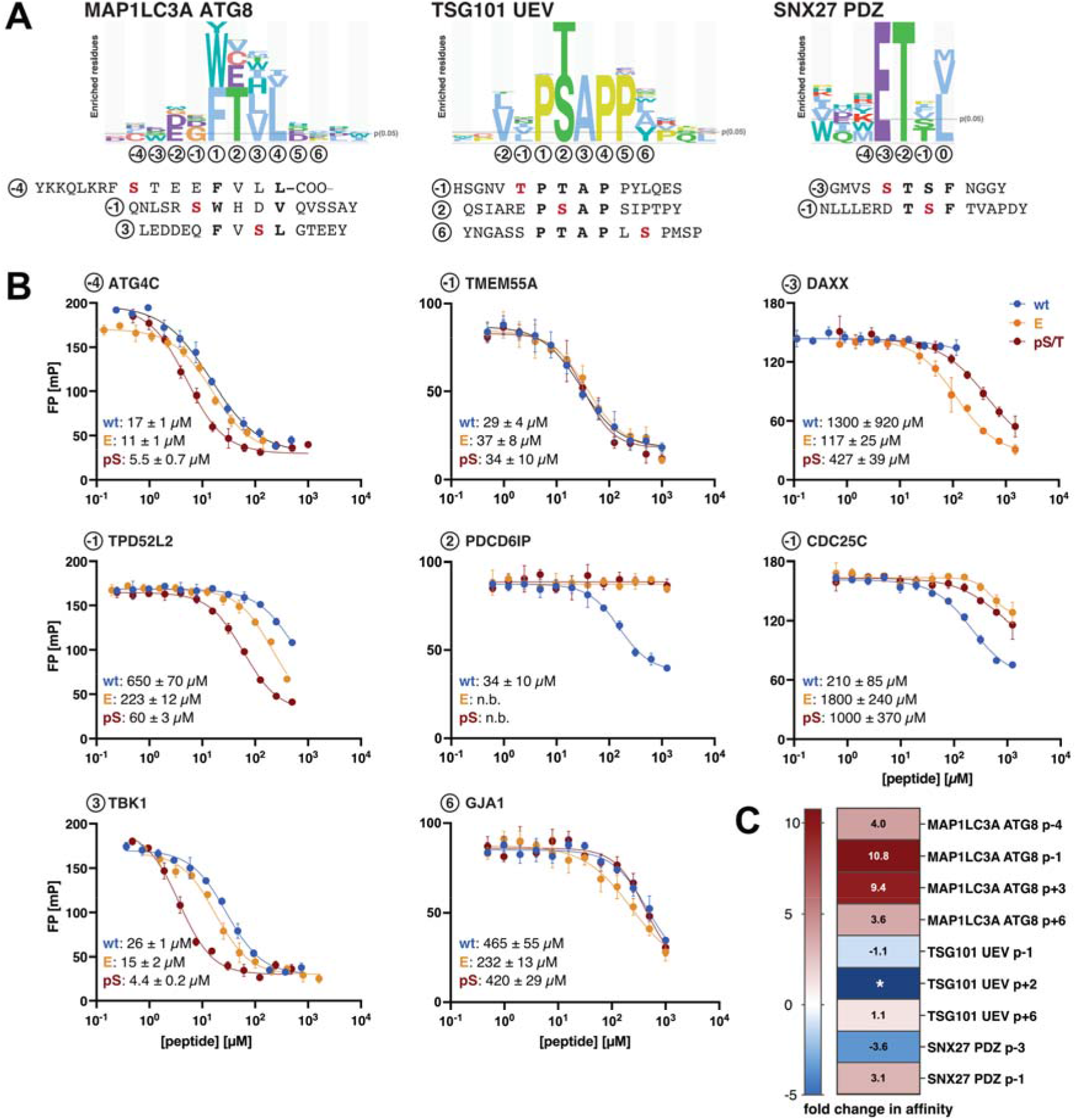
Preferences in binding motifs for serine/threonine or aspartate/glutamate are respectively indicative for negative or positive phospho-modulation. **A**: Binding motifs for MAP1LC3A ATG8, TSG101 UEV domain generated by selections with the HD2 library (20) and the C-terminal PDZ binding motif of the SNX27 PDZ domain. Peptides and phosphosites tested by affinity measurements are indicated and aligned to the motif. **B**: Displacement curves of the three different bait domains and the indicated peptides using wild-type, phosphomimetic and phosphorylated peptide. IC50 values of the different peptides are indicated in μM. **C**: Fold changes in affinity (wild-type vs. phospho-peptides) with blue indicating negative and red indicating positive phospho-modulation of the tested interactions. * The exact fold change for the TSG101 UEV domain - PDCD6IP peptide interaction could not be calculated as the fitting of the displacement curve of the phosphorylated peptide did not provide a IC50 value due to the weak affinity.

The binding of the TSG101 UEV domain to the wild-type, phosphomimetic and phosphorylated PDCD6IP peptide showed that both the phosphomimetic mutation and threonine phosphorylation at the p+2 position abolished binding completely (**Figure 4B, C**). Since serine/threonine at the p+2 position are part of the core P[S/T]AP motif (29), it appears reasonable that phosphorylation would impact binding appreciably. In contrast, modification of the flanking residues at p-1 or p+5 positions of the TMEM55 and GJA1 peptides had no effect. Comparison with the motif from the ProP-PD display supports this observation as neither phosphorylatable (serine/threonine), acidic (aspartate/glutamate) nor basic (arginine/lysine) are enriched at those positions (**Figure 4A**). For the SNX27 PDZ domain, which binds reportedly to a class I PDZ binding motif (30), we found that the affinity was enhanced by phosphorylation at position p-1 and negatively affected by p-3 phosphorylation (**Figure 4B,C**), as indicated by the phage display and reflected by a strong preference for either acidic residues (p-3) or serine/threonine (p-1) at the respective positions (**Figure 4A**). To summarise, we find that it is to some extent possible to postulate the effects of serine/threonine phosphorylation based on consensus motifs generated based on results from selections against a wild-type ProP-PD library but that the phosphomimetic approach provides additional information.

### 2.6 Linking readers to writers and erasers on large scale

Of the 252 high-confidence and prioritised phosphosite interactions identified here, 157 were found to be associated with a kinase (62%) with the vast majority (85%) being represented by mitogen-activated protein kinases (MAPKs) and cyclin-dependent kinases (CDKs), similar to their association with phosphosites in the original library (78%). For most of the phosphosites several kinases are predicted, in which the [ST]P-modifying proline-directed kinases MAPK and CDK targets vastly overlap (MAPK + CDK: 128; CDK only: 6; other: 23) (**Figure 5A**, **Table S1**). In turn, phosphatases could be annotated for 9 out of the 252 phosphosites (3.6%), which represents an increase in percentage in comparison to 0.5% in the PM_HD2 library design (**Figure 1C**). For four cases, we found phospho-switches with information on both the kinases and phosphatases that modify the position. Two examples are the binding of the KPNA4 ARM to the CDC25A114-127 peptide where S116 phosphorylation causes a switch in the pocket preference of KPNA4 ARM domain (**Figure 5A; Table 1**) and the binding of the CREBBP NCBD domain to the STAT1_723-728_ peptide (**Figure 5A,B**; **Table S3**) which is inhibited by the phosphomimetic E727. However, for most cases, there is only information on the predicted kinases. For example, we identified a putative phospho-modulated interaction between the N-terminal domain (NTD) of clathrin and the C-terminal peptide of Hepatoma-upregulated protein (HURP), which was suggested by the display to be positively modulated by phosphorylation at S839 (**Table S3**) by a proline-directed kinase (MAPKs or CDK1) (**Figure 5A, C**).

**Figure 5.**
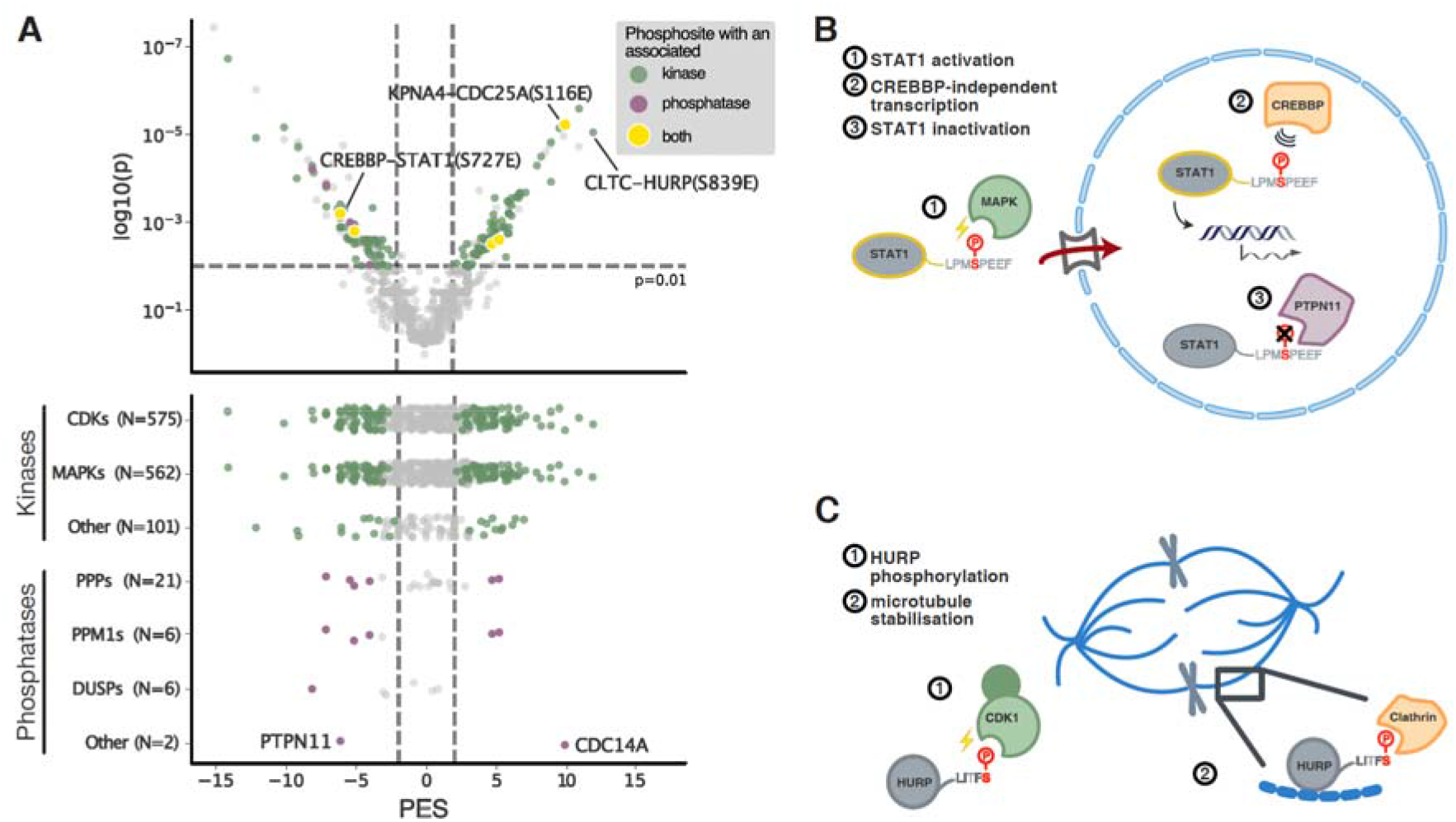
Annotated kinases and phosphatase of the high-confidence and prioritised phosphosite interactions. **A**: **Top**: Kinases (green) and phosphatases (purple) associated with the high-confidence and prioritised phosphosite-interactions. Yellow indicates cases in which both kinase and phosphatase are known for the phosphosite. **Bottom**: Associated kinases and phosphatases of high-confidence phosphosite-interactions across different priorisiation of their effect on binding (grey - insignificant, coloured - significant as judged by the PES and the p-value of the Mann-Whitney test). **B**: Phosphorylation of STAT1 by MAPKs at S727 results in its activation as transcriptional regulator in response to inflammatory stimuli (31,32). The interaction of STAT1 with CREBBP NCBD is inhibited (33), which points at CREBBP-independent transcription upon S727 phosphorylation. Dephosphorylation of S727 is mediated by PTPN11 (34). **C**: Phosphorylation of HURP at S839 by CDK1 enables potentially its interaction with clathrin and together they may contribute to the microtubule stability during mitosis.

### 2.7 Phosphomimetic ProP-PD unravels a phospho-modulated clathrin binding motif in HURP that is necessary for normal mitotic progression

We hypothesised that the interaction between HURP and clathrin can be involved in the regulation of microtubule dynamics at the mitotic spindle as both proteins are reported regulators (35–37). Further, HURP is highly phosphorylated during mitosis suggesting that the proline-directed phosphorylation is a CDK target (35,38,39), which prompted us to investigate the interaction in this cell cycle stage (**Figure 5D**). First, we found that phosphorylation of S839 on the HURP peptide results in a 20-fold increase in affinity, as demonstrated by both FP and ITC experiments (**Figure 6A and B**). To our knowledge, the clathrin NTD has previously only been suggested as a phospho-binder with no structural details of the interaction (40). The clathrin NTD has four reported binding sites to accommodate its adaptor proteins (41), and the identified ligand could potentially bind to one or more of them. To shed light on the structural details of the complex we co-crystallised a longer pS839 HURP_826-846_ peptide with the clathrin NTD. The peptide binds preferentially to the adaptor binding pocket 1 on the clathrin NTD, accommodating the core LIxF motif. although a peptide was also found bound to pocket 3 in one of the monomers. The phosphate group itself is clamped between the basic residues R64 and K96, with the residues in 4.2 Å and 2.7 Å distance respectively, forming salt bridges with the phosphate group (**Figure 6D**). Comparison with the ELM motif (**Figure 6E**) allows good rationalisation for the preference of acidic residues at the p+5 position and hence the phosphorylated serine at the same position in the HURP peptide. Superimposition with previously reported structures of the clathrin NTD (**Figure S6A**) with peptides shows that the acidic residue on the bound peptide assumes a similar, but not identical orientation as the phosphorylated serine in the HURP peptide (**Figure S6B**).

**Figure 6.**
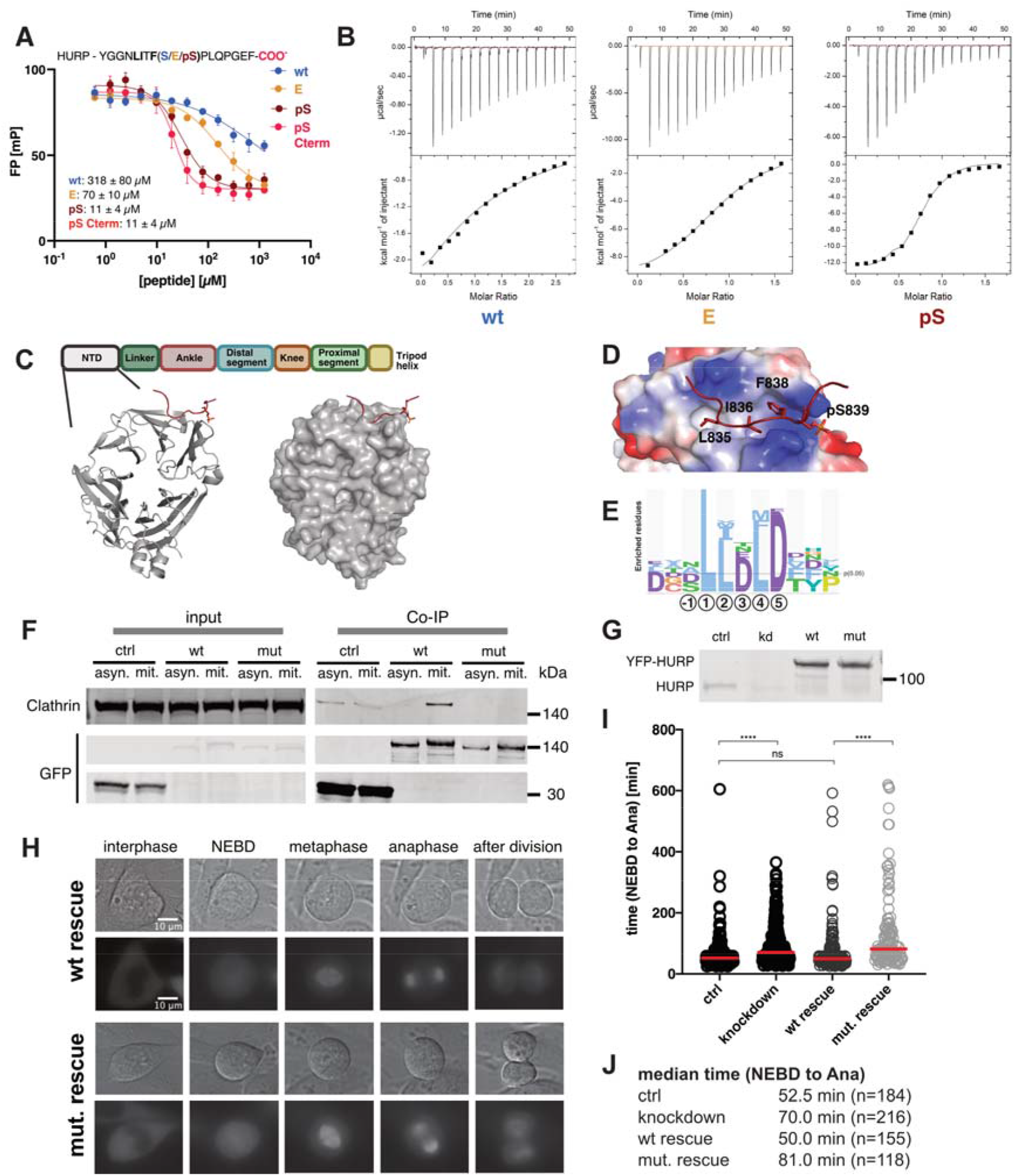
Clathrin binds to a phospho-modulated motif in the HURP C-terminus, which is required for HURP mitotic function. **A:** Displacement curves of the FP experiment using clathrin NTD and the wild-type, phosphomimetic and phosphorylated HURP peptides. IC50 values are indicated in IC50 values of the different peptides are indicated in μM. **B:** ITC experiments of the clathrin NTD and wild-type, phosphomimetic and phosphorylated HURP peptides. **C:** Crystal structure of clathrin NTD with the phosphorylated HURP peptide (PDB: 7ZX4). **D:** Electrostatic representation of the domain bound to the HURP peptide. The phosphate group of Ser839 is complex by two basic residues R64 and K96 **E:** ELM logo of clathrin NTD. **F:** Co-IP of clathrin with Venus-tagged wild-type and mutant HURP constructs in asynchronous and mitotic HeLa cells. **G:** Knockdown of endogenous HURP and inducible expression of RNAi-resistant YFP-tagged wt and mutant HURP. **H**: Representative images of wild-type and mutant HURP HeLa cells in the different mitotic stages. **I-J:** RNAi rescue experiments (n=3), which included the knockdown of endogenous HURP and re-expressing either RNAi resistant wild-type or mutant HURP. In order to assess mitotic behaviour of the cells, the time required of control, HURP knockdown, wild-type HURP and mutant HURP HeLa cells to progress through mitosis (NEBD-anaphase) was established using time-lapse microscopy. Every circle represents one analysed cell. Mann-Whitney test was performed in order to assess significant differences in the median time required for mitotic progression (****: p-value < 0.0001; ns: non-significant)

In order to assess the biological relevance of the discovered clathrin-HURP interaction, we aimed to validate the binding in context of the full-length proteins in a cellular setting. Since S839 is a putative CDK1 site and since HURP is reportedly phosphorylated in mitosis, we focused on this cell cycle stage (38,39). To this end, we generated HeLa cells that stably express inducible Venus-tagged wild-type (HURP_WT_) and mutant (HURP_MUT_) HURP where the last 12 amino acids are deleted. Indeed, we confirmed the interaction between HURP_WT_ and clathrin in mitotic HeLa cells and the interaction was abolished in HURP_MUT_ (**Figure 6F**). In contrast, the co-immunoprecipitation of HURP_WT_ by clathrin failed when it was performed using asynchronous cells, thus supporting that the interaction is restricted to mitosis, when CDK activity is high (**Figure 6F**). Taken together, we confirm the essentiality of the C-terminal motif of HURP for its interaction with clathrin and the interaction to occur specifically in mitosis.

Following these results, we assessed the functionality of the clathrin binding motif for the mitotic function of HURP. It has previously been reported that knockdown of HURP results in a mitotic delay, i.e. in a prolonged time required to progress from nuclear envelope breakdown (NEBD) to anaphase (Ana) (35). Similarly, we observed a delay in mitotic progression when we knocked down endogenous HURP (**Figure 6G**) by RNA interference (RNAi) (median time (knockdown) = 70 min, median time (control) = 53 min); **Figure 6H-J**). To test if HURP interaction with clathrin is necessary for normal mitotic progression, we subsequently induced the expression of RNAi resistant HURP_WT_ and HURP_MUT_. Both proteins displayed the same localisation and associated with the spindle during mitosis. However, while HURP_WT_ efficiently rescued the RNAi of endogenous HURP, HURP_MUT_ did not. In conclusion the association of HURP and clathrin through the C-terminally motif is required for the mitotic function of HURP.

## 3. Discussion

In this study we developed phosphomimetic ProP-PD as a general approach to identify phospho-modulated motif-based interactions. We identified 252 putative phospho-modulated interactions by screening 52 protein domains. Of those, 126 were observed to be enabling and 126 as disabling. We validated the classifications by affinity measurements of 32 wildtype, phosphomimetic and phospho-peptide triplets. We found that phosphomimetic serine/threonine to glutamic acid mutations mimicked the effect of phosphorylation for phosphomodulated interactions, although in most cases with less effect on binding. We further established the PES as a suitable metric for the evaluation of phosphomimetic ProP-PD results. Phosphomimetic ProP-PD correctly identified 77% of the tested interactions in terms of preferential binding to either wild-type or phospho-peptides. This highlights the potential of phosphomimetic ProP-PD to simultaneously provide information on novel interaction partners and on the phospho-modulation of binding.

An alternative way to gain insight into the phospho-modulation of SLiM-based interactions is to analyse the results from conventional ProP-PD selections, and combine the motif-information with the identified binding sites, as previously shown for KPNA4 (20). Phosphorylation of motif-positions that are strictly populated by serine/threonine will most likely have a negative effects on binding. Phosphorylation of serine/threonine that occurs on motif-positions for which there is a preference for acid residues will most likely improve the affinity of the interaction. The motif information can hence aid in the evaluation of the phosphomimetic ProP-PD results by providing an additional level of confidence in a phosphosite suggested by the display. A main advantage of the phosphomimetic ProP-PD is that it opens for the identification of interactions that are enabled by the phosphomimetic mutations and that are not captured using a library of wild-type sequences due to the low affinity of the interaction of an unphosphorylated peptide. The conservative library design of the PM_HD2 library ensures in addition that the phosphosites tested are of high-confidence with respect to their localisation (≥ 0.75 localisation probablity) and functionality (≥ 0.45 phosphosite functional score). This enables the discovery of biologically relevant phosphosites with impact on binding interfaces.

By combining the results of the interaction screening with information on associated kinases and phosphatases, we provide novel insights into the system of phospho-writer, reader and eraser, which can place the interaction in a certain biological setting and in this fashion facilitate potential follow-up studies. One example involves STAT1, the activity of which is heavily regulated by phosphorylation. For instance, S727 phosphorylation has been shown to be required for its activation upon interferon-signalling (31,42). STAT1 cooperates with the CREBBP to regulate transcription (43,44). Screening with PM_HD2 suggested a disabling effect of the phosphomimetic E727, which is consistent with the report of S727 phosphorylation disturbing the interaction with CREBBP (33). It can hence be envisioned that S727 phosphorylation acts, besides its role in STAT1 activation, as a regulatory mechanism to induce transcription independent of CREBBP binding.

The phoshomimetic ProP-PD results revealed moreover a phospho-dependency of the clathrin-HURP peptide interaction. The role of clathrin is well established in endocytosis, but has also been extended to mitosis, during which clathrin plays an essential role in microtubule stability (36,37). Clathrin classically interacts with its adaptors via clathrin adaptor motifs, such as the LIxF motif (20,41). HURP, in turn, is a microtubules binding protein, the knockdown of which results in a mitotic delay (35). The predicted kinase for the phosphosite is CDK, which contextualises the two proteins to a shared biological setting. We showed that this interaction is positively regulated by phosphorylation on a site adjacent to the LIxF motif and provide a structural explanation for its phospho-dependency. We further confirmed the interaction in mitotic HeLa cells and showed that it is required for the mitotic function of HURP. This strengthens the notion that phosphomimetic ProP-PD can delineate interactions with functional relevance in the cell while providing mechanistic details on the amino acid level, including binding motifs and putative phospho-modulation.

Phosphomimetic ProP-PD complements other methods for finding phosphomodulated interactions such as for example using pre-phosphorylated phage libraries (24,45), phospho-peptide pull down coupled to mass-spectrometry (46), yeast-two-hybrid in presence of kinases (15), or using an expanded genetic code (47). The advantage of the phosphomimetic ProP-PD lies within the detailed information on the sites being tested and the scalability of the approach. Of note, screening obligate phospho-binders with the PM_HD2 library was not successful, which pinpoints the limitation of the approach. This limitation applies to all studies using phosphomimetics, such as cell-based studies in which it is frequently used as a proxy for the phosphorylated protein due to the ease of introducing a mutation.

To conclude, we present here phosphomimetic ProP-PD not only as an opportunity to screen for potential phospho-modulated interactions, but also establish a new method for analysing mutational phage display data. The evaluation and classification of the phage display data by the PES to deduct statistically significant binding preferences to either wildtype or mutant peptide can easily be envisioned to be extended to other applications, such as disease-associated mutations. Changes in the interactome by mutations or modifications can be sampled on large-scale due to the feasibility of ProP-PD to screen thousands of different wild-type/mutant peptide pairs.

## 4. Material & methods

### Peptides

Peptides were obtained either from Glbiochem (China) or Genecust (France) and dissolved in dimethyl sulphoxide (DMSO; for FITC-labelled peptides) or 50 mM sodium phosphate buffer pH 7.4 (unlabelled peptides) if not indicated otherwise (**Table S8**). Additionally, peptides were modified if needed to contain a C- or N-terminal tyrosine to allow for concentration determination at 280 nm.

### Protein expression and purification

*Escherichia coli BL21*-Gold(DE3) bacteria were used as expression system. Bacteria were grown to OD_600_ = 0.7, protein expression was induced with 1 mM isopropyl □-D-1 thiogalactopyranoside (IPTG) for 18 h. The bacteria were pelleted for 15 min at 4,000 xg at 4 °C and stored at −20 °C. For purification, pellets were resuspended in lysis buffer (phosphate buffered saline (PBS) pH 7.4, 5 mg/mL MgCl_2_, 0.05% Triton-X, lysozyme, 10 μg/mL DNAse and cOmplete™ EDTA-free Protease Inhibitor Cocktail (Roche)), incubated for 1 h at 4 °C and sonicated before spin down at 16,000 xg at 4 °C. For phage selections, proteins were expressed as GST- or MBP-tagged fusion proteins (**Table S3**) and purified from the supernatant with glutathione (GSH) beads (Cytiva) or Ni^2+^ Excel beads (Cytiva), respectively.

For FP, ITC and crystallisation, the tag was removed with the appropriate protease either on beads (0.1 mg/mL thrombin (Cytiva) in PBS, 1 mM dithiothreitol (DTT)) or by dialysis in cleavage buffer (Human Rhinovirus (HRV) 3C protease: 50 mM Tris HCl pH 7.4, 150 mM NaCl, 1 mM β-mercaptoethanol), both procedures were overnight. In case of HRV protease cleavage, the tag was separated from the target protein domain by reverse ion metal affinity chromatography using Ni^2+^ beads for purification, whereas thrombin was removed using a benzamidine column (Cytiva). Affinity measurements for SNX27 PDZ were performed on the MBP-tagged construct, as well as PIN1 WW domain on the GST-tagged construct. For FP and ITC, proteins were dialysed in 50 mM sodium phosphate buffer pH 7.4, 1 mM DTT. For crystallisation of the NTD of clathrin, the protein was further purified by a S200 Sepharose column (Cytiva) using 20 mM Tris HCl pH 7.4, 150 mM NaCl and 1 mM DTT.

### Design of the PM_HD2 library

The PM_HD2 library was designed on the bases of our previously published HD2 library (20) by systematically mutating serines and threonines identified in a previously published MS-defined phospho-proteome (23) to glutamate. First a list of phosphosites was built (**Table S2A**). To keep the scale of the library to ~92,000 peptides only serine and threonine phosphosites were considered and they were filtered by localization probability (≥ 0.75) and functional score (≥ 0.4524). For proteins where none of its phosphosite surpassed the cut-offs, the single phosphosite with the highest functional score was also kept. A list of proteins of interest was considered to rescue some phosphosites back, these proteins of interest included KPNA4, a list of curated switches and ELM instances containing serine or threonine phosphosites (**Table S2B**) and a list of curated NLS/NES (**Table S2C**). Finally, this list of phosphosites was mapped onto the HD2 library design allowing only a single mutation at a time if multiple phosphosites occurred in the same peptide, and a wild type and a glutamate version of each peptide were reverse translated into oligonucleotides using optimised codons for *Escherichia coli* expression avoiding the formation of SmaI restriction sites (**Table S1**).

Predicted kinases for all phosphosites in the PM_HD2 library were obtained from Ochoa et al., 2020 (23) and phosphatases were obtained from DEPOD (59).

### Generation of the PM_HD2 library

The PM_HD2 library was, as previously described, generated on the p8 protein of the M13 bacteriophage by first amplifying the custom oligonucleotide array, annealing the amplified array to the single-stranded phagemid which was followed by amplification of the phagemid (Kunkel reaction). After that, the library is transformed by electroporation into SS320 bacteria (20,48).

1. The custom oligonucleotide array (1 μL) was amplified with primers for internal p8 coat protein display and Phusion High-Fidelity PCR Master Mix (Thermo Scientific).
2. Amplification was verified by running the PCR product on a 2% agarose gel and the PCR products pooled and cleaned up with QIAquick Nucleotide Removal Kit (Qiagen). The DNA content was quantified with Quant-iT PicoGreen dsDNA Assay Kit.
3. The dsDNA product was denatured for 10 min at 98 °C. Next, the amplified custom library (0.3 μg) was phosphorylated with the T4 polynucleotide kinase (10 units) for 1 h at 37 °C. To the phosphorylated oligonucleotides, 10 μg dU-ssDNA of the p8 phagemid were added (ratio 3:1) and the sample incubated for annealing 3 min at 90 °C, 3 min at 50 °C and last 5 min at 20 °C.
4. For amplification of the phagemid, 30 units of T7 DNA polymerase and 30 Weiss units of the T4 DNA ligase were used and the sample incubated for 20 h at RT.
5. Wild-type phagemid was removed by SmaI digestion (5 μL) for 30 min at 37 °C. The CCC-dsDNA was then purified QIAquick PCR Purification Kit and success of the Kunkel reaction verified by running the sample on a 2% agarose gel.
6. The library was transformed into electrocompetent SS320 bacteria (150 μL), which were pre-infected with M13KO5 helper phages, and using a XX electroporator. The sample was immediately resuspended in 1 mL pre-warmed SOC medium, transferred to additional 9 mL medium and incubated for 30 min at 37 °C at 220 rpm. The 10 mL were transferred after that to a 500 mL culture containing 30 μg/mL kanamycin and 100 μg/mL carbenicillin and the phage library grown o.n. at 37 °C at 220 rpm.
7. The bacteria were pelleted by centrifugation at 4,500 xg for 15 min at 4 °C, the phages precipitated by addition of one-fourth of the volume of 20% PEG-800 and 0.4 M NaCl to the supernatant, incubation for 10 min on ice and subsequent spin at 16,000 g for 10 min at 4 °C. The pellet was resuspended in 20 mL 0.05% Tween-20 in PBS, ultrapure glycerol added (10%) and the library stored at −20 °C.

### Phage display selections

Selections using the internal phosphomimetic phage library were performed as described previously with minor modification to the original protocol. The corresponding tag protein (either GST or MBP) were first used during negative selections. Selections were run for four days.

1. Bait and tag proteins (10-15 μg in 100 μL PBS) were immobilised in 96-well MaxiSorb plate (NUNC) o.n. at 4 °C.
2. Escherichia coli OmniMax bacteria were grown in 2YT media supplemented with tetracycline (100 μg/mL) at 37 °C and 200 rpm.
3. Plates with immobilised bait / tag proteins were blocked with 0.5% BSA in PBS for 1 h at 4 °C.
4. Phage library (10 μL/well) was prepared by diluting 10x with PBS and adding one-fourth of the volume of PEG/NaCl. For phage precipitation, prepared library was incubated for 10 min on ice followed by centrifugation for 10 min at 10,000 xg at 4 °C. Phages were resuspended in 0.5% BSA, 0.05% Tween in PBS.
5. Plates with tag proteins were washed 4x with 0.05% Tween in PBS, *naïve* phage library was added to each well (100 μL/well) and the plates incubated for 1 h at 4 °C.
6. Plates with bait protein were washed 4x with 0.05% Tween in PBS, non-bound phages were transferred from the tag protein plate and the plates incubated for 1 h at 4 °C.
7. Plates were again washed 5x with 0.05% Tween in PBS and bound phages were eluted with log-phase OmniMax bacteria (100 μL/well) by incubating for 30 min at 37 °C at 200 rpm.
8. M13K07 helper phage (10^12^ cfu/mL) were added to each well and the plates incubated for 45 min at 37 °C at 200 rpm.
9. Amplification took place o.n. by adding the eluted phages to 1.1 mL 2YT media supplemented with kanamycin (25 μg/mL), carbenicillin (50 μg/mL) and 0.3 μM IPTG in a 96 deep well plate and incubating o.n. at 37 °C at 200 rpm.
10. Bacteria were pelleted by centrifugation for 15 min at 1700 xg at 4°C. The phage supernatant was pH-adjusted by addition of 10x PBS and remaining bacteria heat-inactivated by incubating for 10 min at 60 °C. Selections were repeated as described from (5).

### Phage Pool ELISA

1. Phage Pool ELISA on selection days 1-4 was performed as previously described (20,48). Briefly, bait domains and control proteins (5-7.5 μg of protein per well in 50 μL PBS) were immobilised in a 384-well MaxiSorb plate (NUNC) o.n. at 4 °C while shaking.
2. Plates with immobilised protein were blocked with 0.5% BSA in PBS and incubated for 1 h at 4 °C.
3. The blocking solution was removed and the phage pools from the various selection days were added to the respective bait domains and their control protein, and the plates incubated for 1-2 h at 4 °C while shaking.
4. The plates were washed 5x with 0.05% Tween in PBS, 100 μL of anti-M13 bacteriophage mouse antibody (Sino Biological Inc, 1:5000) were added to each well and the plates incubated for 1 h at 4 °C while shaking.
5. The antibody solution was washed off by washing the plates 4x with 0.05% Tween in PBS and 1x with PBS. To each well, 40 μL of TMB (3,3’,5,5’-Tetramethylbenzidine) substrate were added and the enzymatic reaction subsequently stopped with 40 μL of 0.6 M H_2_SO_4_. The absorbance of bait protein domain and control protein at 450 nm was subsequently compared. The enrichment of phages was considered specific for the bait protein domain when the ratio at 450 nm was above 2.

### Next generation sequencing (NGS)

Based on the results from Phage Pool ELISAs, samples from selection day three and four were selected for NGS, as described before (20).

1. Peptide coding regions were amplified and dual-barcoded by PCR. 5 μL of the enriched phage pools (50 μL reactions) and a high-fidelity phusion polymerase (Roche) were used for the PCR reaction. Amplification was confirmed by running the PCR product on 2% agarose gel with a 50 bp ladder.
2. PCR products (25 μL) were purified with 25 μL of Mag-bind Total Pure NGS beads and normalised in the same step by taking 10 μL of each reaction after resuspension (Qiagen Elution buffer: 10 mM Tris-Cl and 0.1 mM EDTA (pH 8.5)) from the beads.
3. The samples were cleaned up with a PCR purification kit (Qiagen), subsequently run on a 2% agarose gel and bands of the correct size (ca. 200 bp) were purified with the QIAquick Gel extraction Kit (Qiagen). From the columns, samples were eluted with 30 μL TE (10□mM Tris-HCl, 1□mM EDTA. pH 7.5) buffer and quantified with Quant-iT PicoGreen dsDNA Assay Kit.
4. Samples were sequenced on MiSeq (MSC 2.5.0.5/RTA 1.18.54) with a 151nt(Read1) setup using ‘Version3’ chemistry. The Bcl to FastQ conversion was performed using bcl2fastq_v2.20.0.422 from the CASAVA software suite.

### Sequencing analysis

The processing of sequencing data was performed as described in Ali *et al*., 2020 (21). In brief: Pooled results from ~500 selection experiments were demultiplexed by identifying unique combinations of 5’ and 3’ barcodes, sequences with an average quality score of 20 or more were conserved and their adaptors were trimmed and DNA sequences translated using in house custom Python scripts. A table for each experiment was built containing peptide sequences associated to their sequencing counts.

### Library coverage and quality

The completeness of the PM_HD2 library was evaluated by sequencing multiple non-challenged *naïve* library aliquots. The observed coverage of the library was evaluated and the maximum estimated coverage was determined (**Figure S1A**) as described in Benz *et al*., 2022 (20). Finally, the counts distribution per peptide was evaluated (**Figure S1B**) and the percentage of wild type-mutant peptide pairs and their ratios distributions were assessed (**Figure 1D**).

### Selection results analysis

After building tables *per* experiment as described in the “Sequencing analysis” section peptide sequences were analysed as described in *Benz et al*., 2022 (20) with some modifications regarding evaluation of wild-type vs mutant versions of each peptide pair. In brief: Peptides with a read count of 1 and those peptides that did not match the library design were removed and the remaining peptide counts were normalized. Different selection days for the same replica experiment were merged and their normalized counts average was calculated for each peptide.

Different replicates for the same bait were joined together and PepTools was used to annotate all peptides and each peptide’s confidence score was calculated as described previously (20). Finally, a mutation centred analysis was performed by evaluating the selection read counts for all wild type and mutant overlapping peptides for each phosphosite. This analysis included calculating a Mann-Whitney confidence score by comparing the counts from wild type vs mutated peptides, and the calculation of the Phosphomimetic Enrichment Score (PES) for the phosphosite following the equation:

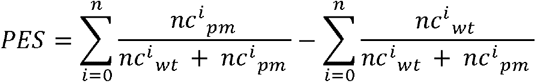

**Equation 1**. The phosphomimetic enrichment score (PES) is calculated on the level of wt and phosphomimetic mutation pair in selection against a given bait. The score includes data for overlapping peptide containing the same wild-type and phosphomimetic pair. The PES is calculated using the normalised sequencing counts for the wild-type (*nc^i^_wt_*) and phosphomimetic (*nc^i^_pm_*) data for all replicates for all peptides. Each wt and phosphomimetic mutation pair will have n datapoints and the bounds of the score are +/- n. A positive score indicates a phosphomimetic enhances binding and a negative score indicates a phosphomimetic inhibits binding.

### FP experiments

FP experiments were performed as described previously (49). Shortly, in order to monitor the binding event, FITC-labelled peptides were used (excitation 495 nm, emission: 535 nm) and the FP signal was recorded in mP on a SpectraMax iD5 Multi-Mode Microplate Reader by measuring the emitted light in two angels. The measurements were performed at room temperature and prepared in non-binding black half area 96-well microplates (Corning^®^) (50 μL/well). For measuring direct binding, i.e. saturation curves, a 1:1 dilution series of protein domain (25 μL/well) was generated in 50 mM sodium phosphate buffer pH 7.4 and the measurement performed with 5 nM FITC-labelled peptide (25 μL/well). The buffer-corrected values were subsequently plotted with PRISM and fitted to the quadratic equation (50).

In order to obtain the K_D_ values of the direct binding event. In order to record displacement curves, a complex of protein domain ([domain] = 1-2 fold over K_D_ of FITC-labelled peptide) and FITC-labelled peptide (10 nM in complex, 5 nM in the measurement) was used. The 1:1 dilution series in this experiment was performed with the unlabelled peptide in 50 mM sodium phosphate buffer pH 7.4, so that each well contained unlabelled peptide (25 μL) at a certain concentration and pre-formed complex (25 μL). The K_D_ values of this indirectly monitored binding event were calculated as described previously (51). All measurements were at least in technical triplicates as well as biological duplicates when indicated.

### ITC experiments

Both peptide and protein domains were dialysed in the same buffer to avoid buffer mismatch causing artefacts (50 mM sodium phosphate buffer pH 7.4, 1 mM DTT). Experiments were performed on an iTC200 instrument (Malvern) by titrating peptides against protein domains in approximately 10-fold excess (16 injections). The provided program on the instrument was used for curve fitting and manual baseline correction was applied if suitable. Technical triplicates were obtained for all measurements, except for the GGA1 VHS domain binding to the RNF11 peptide for which a technical duplicate has been obtained.

### Crystallisation

For crystallisation, Clathrin NTD (19 mg/mL) and HURP pS839 C-terminal peptide (6.4 mM in 50 mM sodium phosphate buffer pH 7.4) were mixed in 1:1 molar ratio. Initially, the crystallisation was attempted by using reported crystallisation condition (50 mM Tris pH-7.5 and 30% PEG 6000) (36). The hanging drop method was applied, correspondingly 2 μL of complex was mixed with 2 μL of reservoir solution of composition 24% PEG-6000, 100 mM Tris-HCl pH 7.5 and pH 8.0. The crystallisation was performed at 293 K and crystals grew within three days. CLTC-NTD complex crystals were cryoprotected in mother liquor supplemented with 20% glycerol and flash frozen in liquid nitrogen.

### X-ray data collection, structure determination and refinement

For the two peptide complexes of CLTC NTD, crystallographic data was collected at 100 K at the beamline ID-23-1 at European synchrotron Radiation Facility (Grenoble, France) and autoindexed and processed using Xia2 (52). The structures were solved by molecular replacement using Phaser (53) and PDB entry 1C9I as search model (54). Structure was refined with phenix.refine and Refmac5 of the Phenix (55) and CCP4 program suites (56), respectively. Manual model building was done in COOT (57). The final structures showed good geometry as analysed by Molprobity (58). The data collection and refinement statistics are given in **Table S11**.

### Co-immunoprecipitation

For Co-IP experiments, HeLa cells expressing either YFP-tagged wild-type HURP, mut. HURP or YFP only were cultured in selection medium (DMEM 10% FBS, 1% Penicillin/streptomycin, 5 μg/mL blasticidin, 0.1 mg/mL hygromycin) at 37 °C, 5% CO_2_ in a humid environment. Cells were synchronised by 24 h incubation with 2.5 mM thymidine and either collected (asynchronous) or released with 200 ng/mL nocodazole (mitotic). Mitotic cells were collected by shake-off after 20 h. Expression was induced for both asynchronous and mitotic cells for 24 h at 5 ng/mL doxycycline. After collection, cells were re-suspended in lysis buffer (100 mM NaCl, 50 mM Tris-HCl pH 7.4, 0.05% NP-40, 1 mM DTT and completed with phosphatase (PhosSTOP (Roche)) and protease (cOmplete EDTA-free (Roche)) inhibitors) and subsequently lysed by sonication. Lysate was cleared by centrifugation at 20000 xg at 4 °C for 45 min. GFP-trap beads (Chromotek) were used for pulldown and incubated with supernatant for 1 h at 4 °C on a rotating wheel. After that, beads were washed (150 mM NaCl, 50 mM Tris-HCl pH 7.4, 0.05% NP-40, 1 mM DTT and completed with phosphatase and protease inhibitors as above) and eluted with 2-fold NuPAGE LDS Sample buffer (Novex). Samples were separated by SDS-PAGE and blotted in transfer buffer (25 mM Tris, 192 mM glycine, 20% ethanol) at 200 mA at 4 °C for 3 h. Primary antibodies (anti-CHC; anti-HURP; anti-GFP) were diluted 1:1000 in 2.5% milk PBST and incubated o.n. at 4 °C. Blots were washed with PBST, incubated with secondary antibodies (IRDye 800CW anti-mouse; IRDye 680RD anti-rabbit) for 1 h at room temperature and washed again. For visualisation, Odyssey CLx (LI-COR) was used.

### RNA interference (RNAi) rescue experiments

For the knockdown experiments, HeLa cell lines expressing wild-type HURP, mut. HURP or YFP only were transfected with either HURP or control siRNA (10 nM final concentration) using Lipofectamine™ RNAiMAX Transfection Reagent (Invitrogen) in OPTIMEM medium (Invitrogen), which was supplemented after 6 h with 10% FBS. After x h, the cells were seeded at 60% in ibidi dishes in selection medium and synchronised for 24 h with 2.5 mM thymidine. The cells were washed to release them from the thymidine block and and medium was changed to L15 medium with 10% FBS and 1 ng/μL doxycycline. The ibidi dishes were mounted for time-lapse microscopy 2 h after induction on a Delta Vision microscope (Leica). Mitotic progression was monitored by taking images every 5 min for 20 h at different areas of the wells and three z-stacks (5 μm apart). Image analysis was done with Deltavision Softworx Software and Fiji.

## Supporting information

Overview of Supplementary information and Supplemental Figures 1-6

## Material table (Table S12)

## Acknowledgement

This work was supported by the grants from the Swedish Research Council (YI: 2020-03380), the Ollie and Elof Ericsson foundation (YI), the Carl Trygger foundation and the Cancer Research UK (CRUK) (NED: Senior Cancer Research Fellowship C68484/A28159). Sequencing was performed by the SNP&SEQ Technology Platform in Stockholm. The facility is part of the National Genomic Infrastructure (NGI) Sweden and Science for Life Laboratory and is also supported by the Swedish Research Council and the Knut and Alice Wallenberg. Work at the Novo Nordisk Foundation Center for Protein Research is supported by grant NNF14CC0001. We thank Dr. Pedro Beltrao and Dr. David Ochoa for useful discussions on the library design. We thank Dr. Sachdev Sidhu for providing the phagemid, and several of the expression constructs.

## Author contribution

LS designed the phage library with support of NED and DO. JK constructed the phage library, purified proteins, performed phage selections, affinity measurements and analysed the data. JK performed and analysed cell-based validations with support of DHG and JN. DB and DD solved the structure. NED and LS processed ProP-PD data. JK, DHG, JN, NED and YI designed experiments. NED and YI conceived the study. JK, LS, NED and YI wrote the paper with input from all authors.

