## Supplementary material for "Large-scale identification of phospho-modulated motif-based protein-protein interactions": Overview of Supplementary information and Supplemental Figures 1-6

**Supplementary information overview**

**SUPPLEMENTAL FIGURES (provided in this document)**

**Figure S1**	Library quality

**Figure S2**	FP binding experiment 14-3-3 theta and PIN1 WW

**Figure S3**	Probe peptide binding

**Figure S4**	Competitive FP experiment for wild-type, phosphomimetic and phosphorylated peptide triplets.

**Figure S5**	ITC data

**Figure S6**	Comparison of clathrin NTD binding to a non-phosphorylated peptide.

**SUPPLEMENTAL TABLES (provided as separate files)**

**Table S1**	PM_HD2 library design

**Table S2**	Phosphosites collection for the PM_HD2 design

**Table S3**	Bait protein domain collection

**Table S4**	Bait protein domain categorisation

**Table S5**	High/medium confidence ProP-PD data

**Table S6**	Ranking of the medium/high confidence-data set for phospho-modulated

binding.

**Table S7**	Comparison with Switches.ELM

**Table S8**	Detailed peptide information

**Table S9**	K_D_ values from the saturation experiments with FP

**Table S10**	ITC parameters

**Table S11**	X-ray statistics

**Table S12**	Materials

**SUPPLEMENTAL FIGURES**

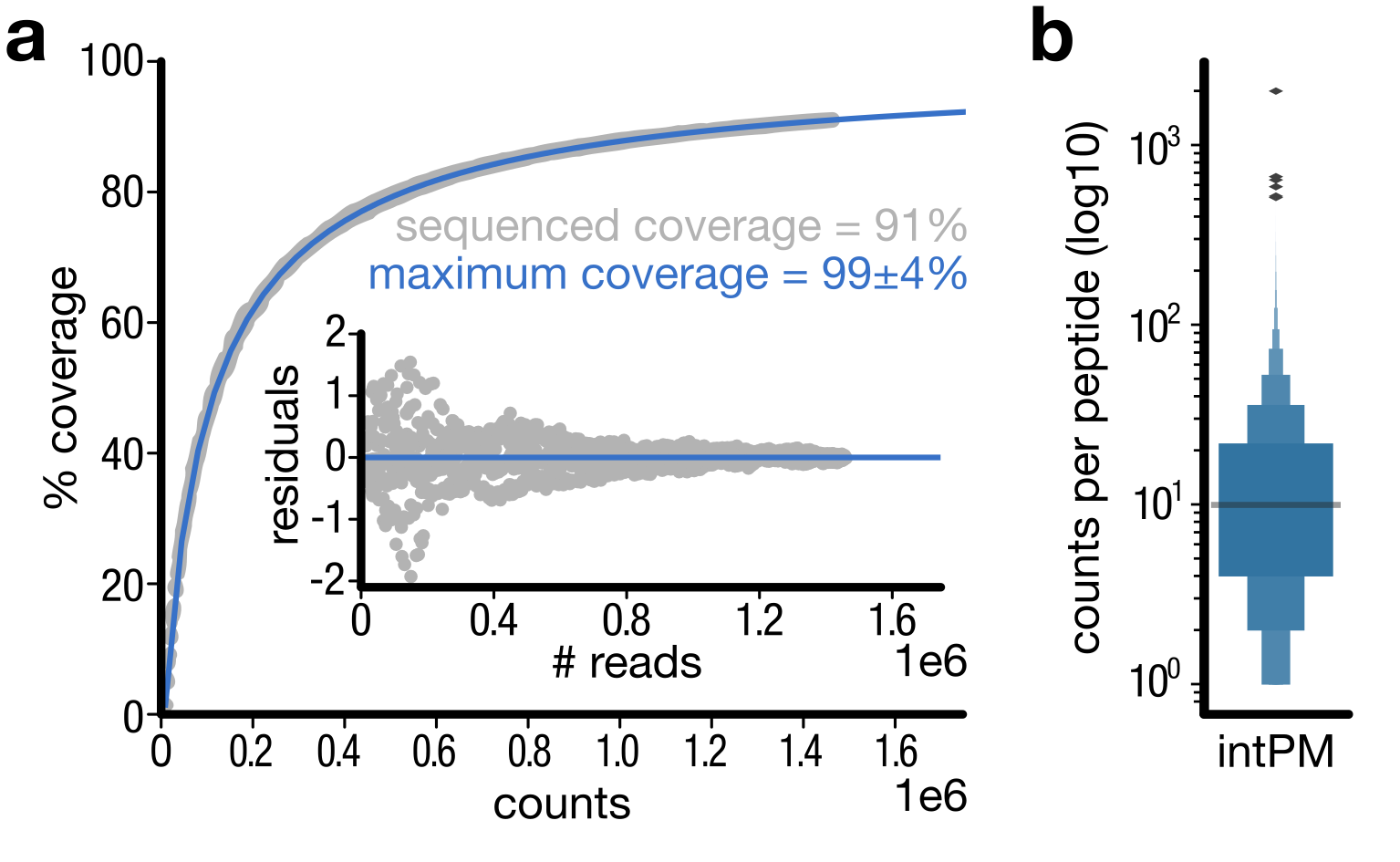

**Figure S1. PM_HD2 library quality A:** Peptide coverage obtained after multiple sequencing of the *naïve* PM_HD2 library. The maximum coverage was calculated by fitting the coverage percent vs. number of reads to a double hyperbolic curve as described in Benz *et al.* 2022. **B:** Counts per peptide distribution in the *naïve* sequencing of the PM_HD2 library.

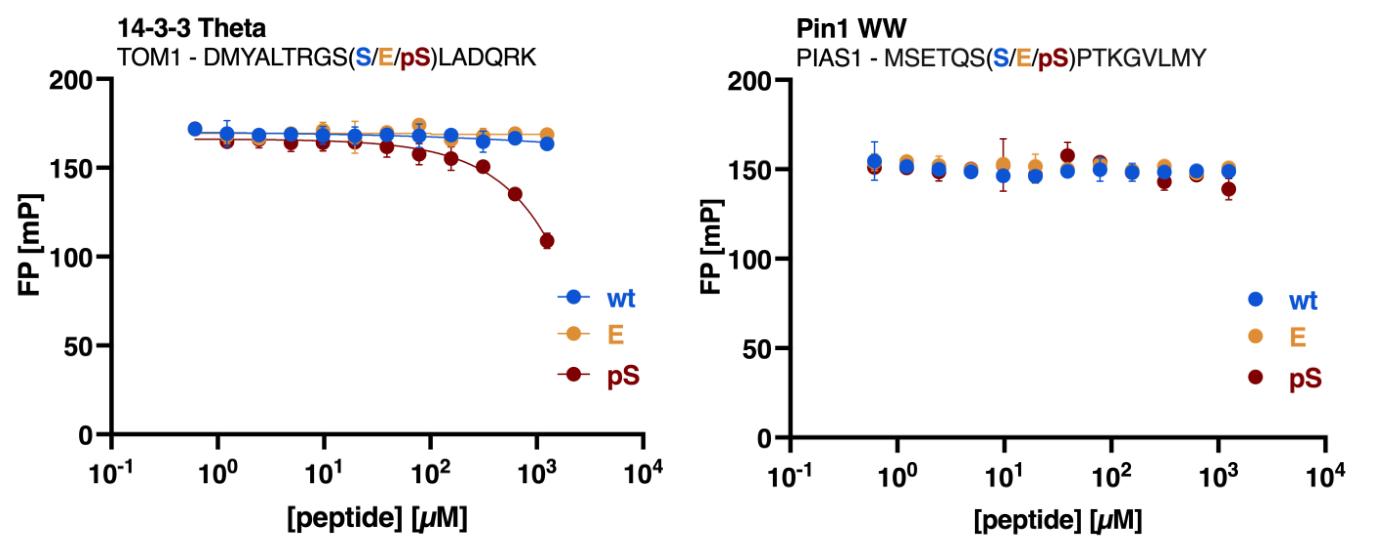

**Figure S2**: Displacement curves of FP experiments of 14-3-3 Theta and PIN1 WW domain with wild-type, phosphomimetic and phosphorylated peptides from TOM1 and PIAS2, respectively.

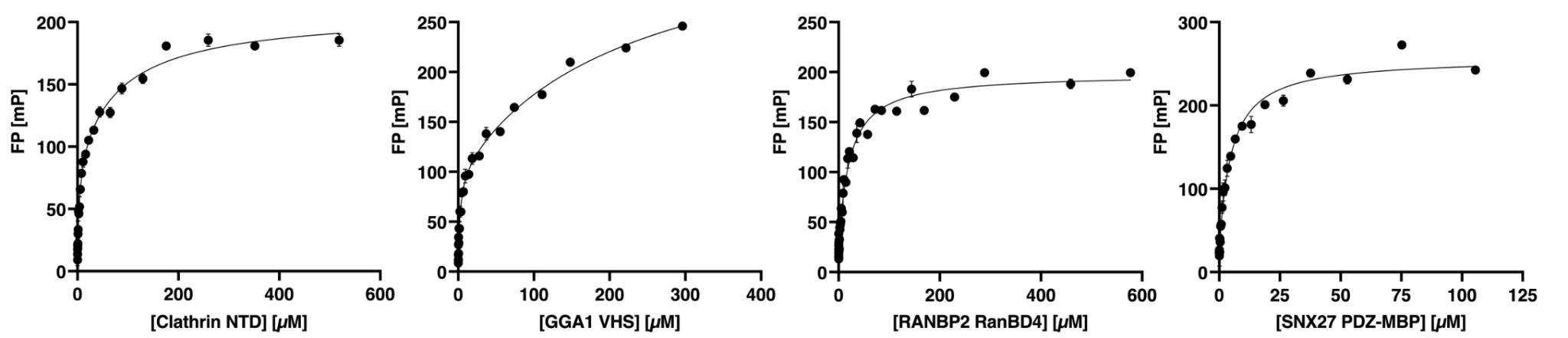

**Figure S3**: Saturation curves of protein domains and FITC-labelled peptides, which have been used in this study but the affinities of which have not previously been reported.

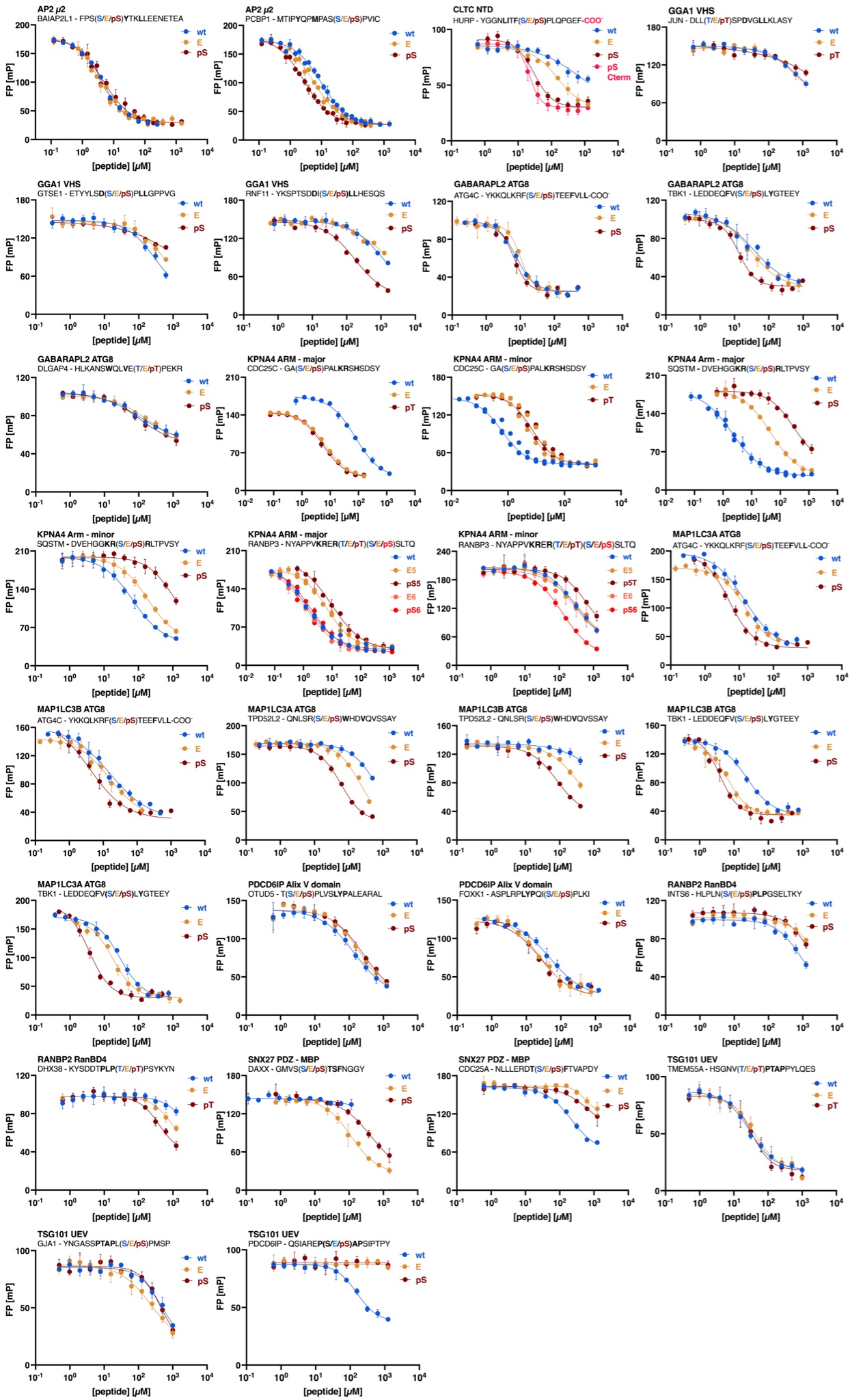

**Figure S4**: Displacement curves of the different protein domains and the peptide triplets (wild-type, phosphomimetic and phosphorylated).

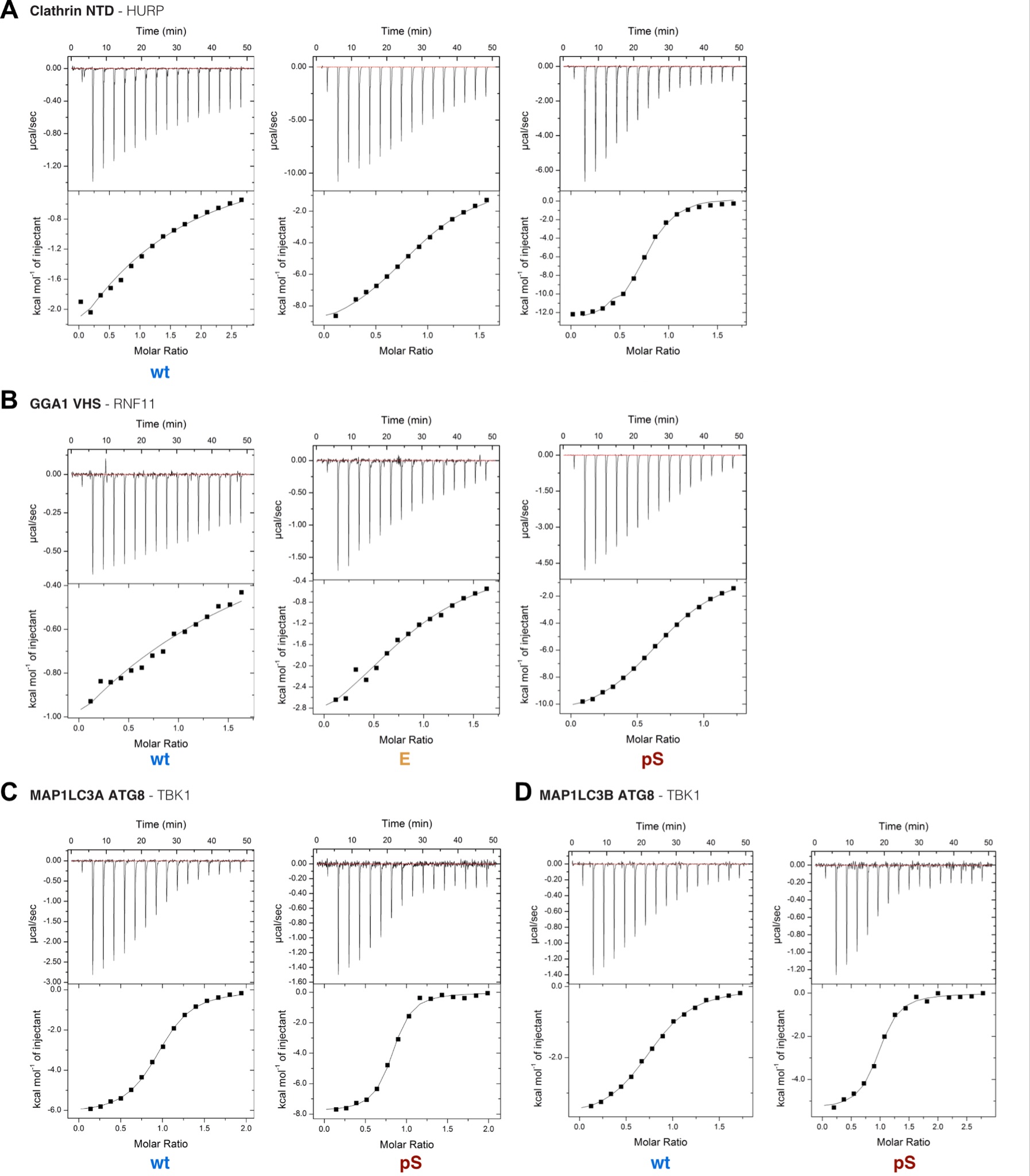

**Figure S5**: ITC curves of clathrin NTD binding to the HURP peptide (wild-type, phosphomimetic and phosphorylated), GGA1 VHS binding to the RNF11 peptide (wild-type, phosphomimetic and phosphorylated) and the ATG8 proteins MAP1LC3A and -B binding to the TBK1 peptide (wild-type and phosphorylated). Measurements were in technical triplicates, except for the GGA1 VHS binding curves which were in duplicates.

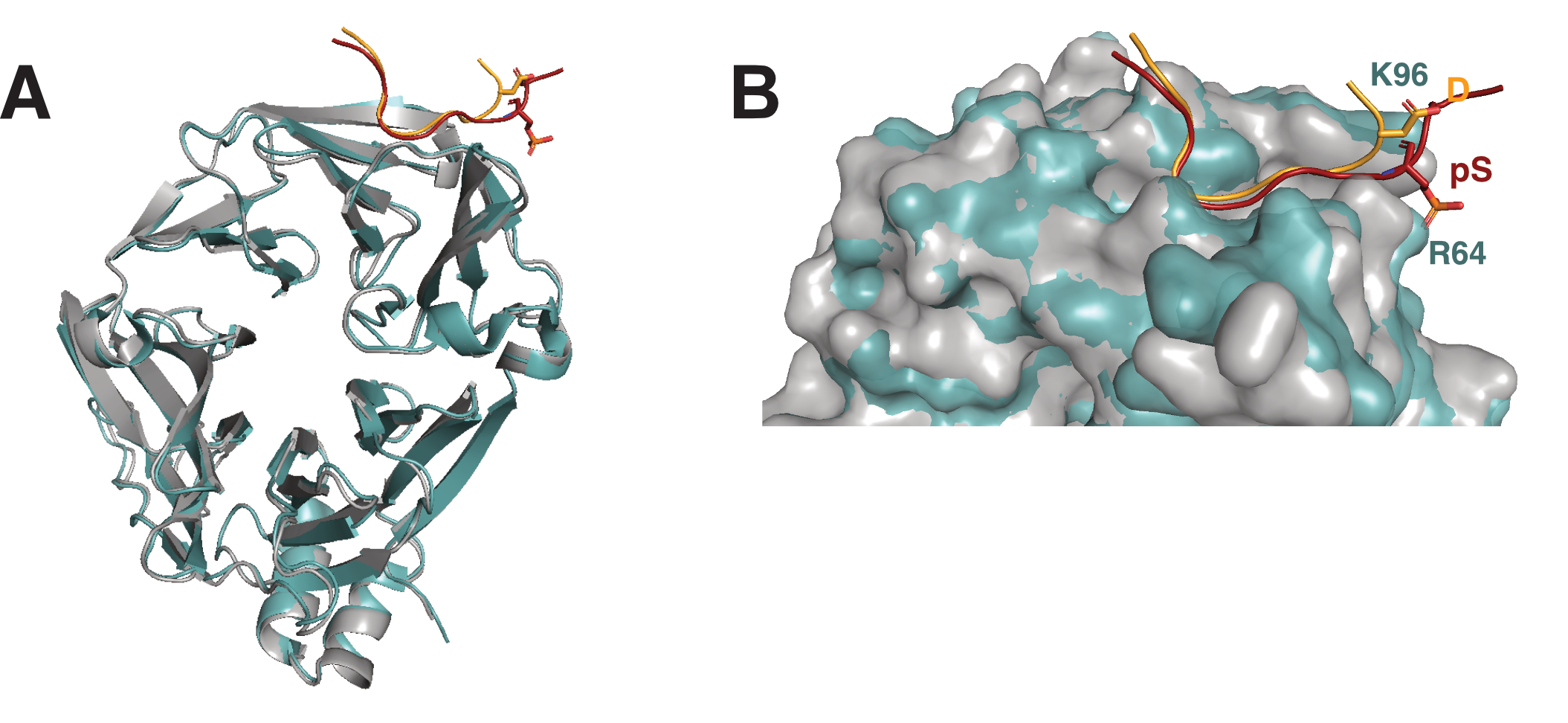

**Figure S6**: Superimposition of clathrin NTD structure bound to non-phosphorylated (PDB: 1C9I) and phosphorylated peptide (this study). A: Domain structure with indicated peptide. B: Comparison of the orientation of the residue at the p+5 position on the peptide, which is either an aspartate (1C9I) or the phosphorylated serine (this study).
