## Supplemental Table 11 for "Large-scale identification of phospho-modulated motif-based protein-protein interactions"

**Supplemental Table 11: Data collection and refinement statistics.** Values given in parentheses are for the highest resolution shell.

|  | **CLTC-HURP pS839** |
| --- | --- |
| **Data Collection** |  |
| Beamline | ESRF ID23-1 |
| Space group | P2_1_ |
| *a*, *b*, *c* (Å), β (°) | 48.8, 94.2, 89.0, 103.8 |
| Wavelength (Å) | 0.8731 |
| Resolution (Å) | 43.20-2.08 (2.12-2.08) |
| Total reflections | 122433 (6291) |
| Unique reflections | 45105 (2302) |
| Multiplicity | 2.7 (2.7) |
| Completeness (%) | 96.0 (97.7) |
| I/σ(*I*) | 6.9 (1.3) |
| Wilson B factor (Å^2^) | 22.4 |
| R*_merge_* | 0.099 (0.390) |
| R*_meas_* | 0.123 (0.475) |
| R*_pim_* | 0.071 (0.268) |
| *CC_1/2_* | 0.909 (0.716) |
| **Refinement** |  |
| No. of reflection in work set | 42817 |
| No. of reflection in free set | 2286 |
| R*_work_* | 0.2061 |
| R*_free_* | 0.2490 |
| Protein-peptide complexes/AU | 2 |
| No. of non-hydrogen atoms/Average B-factor (Å^2^) |  |
| Protein | 5600/30.14 |
| Solvent | 329/31.95 |
| Peptides | 231/37.37 |
| RMS deviations |  |
| bonds (Å) | 0.007 |
| angles (°) | 1.440 |
| Residues in Ramachandran plot regions (%) |  |
| favored | 97.7 |
| allowed | 2.3 |
| outliers | 0.0 |
| PDB-ID | 7ZX4 |
